# High-Performance All-Atom Continuous Constant-pH Molecular Dynamics and Native Asynchronous Replica Exchange in Amber

**DOI:** 10.64898/2026.11.11.766022

**Authors:** Guy W. Dayhoff, Jana Shen

## Abstract

Conformational dynamics of biomolecules, including proteins and nucleic acids, are often coupled to changes in protonation states. All-atom particle-mesh Ewald continuous constant-pH molecular dynamics (PME-CpHMD) enables proton-coupled dynamics, but its routine use has been limited by computational cost. Here we report a second-generation CUDA implementation of PME-CpHMD in Amber, in which the CpHMD-specific nonbonded contributions are evaluated in a separate kernel. Across protein systems ranging from 45 to 1,338 residues, the new implementation increases throughput by 1.5-2 fold relative to the previous fused-kernel implementation and limits the overhead relative to conventional MD to 12–18%. We also implemented the asynchronous pH replica-exchange (pH-REX) algorithm natively in pmemd.cuda.MPI, allowing simulations to run on fewer GPUs than replicas without the overhead of the previous Python wrapper. Furthermore, we derived CpHMD titration parameters for all six titratable residue types (Asp, Glu, His, Cys, Tyr, and Lys) for use with three protein force fields: CHARMM c22/CMAP, Amber ff14sb and ff19sb. The new implementations and parameterization were validated using p*K*_a_ calculations of model peptides and hen egg white lysozyme (HEWL). The HEWL simulations revealed force-field-specific limitations, including overestimation of the p*K*_a_ downshift of H15 and sensitivity of the proton donor assignment in coupled E35/D52 dyad. Comparison with our previous work and the PME-CpHMD simulations in GROMACS with the CHARMM c36m force field demonstrated the robustness and sampling efficiency of pH-REX PME-CpHMD simulations. The new implementation provides a foundation for routine all-atom constant-pH simulations in Amber. The analysis highlights the need for continued development of protein force fields to more accurately balance intramolecular electrostatics and solvation of charged residues.

## 1 Introduction

Protonation state changes are central to many protein processes, including folding,^1^ catalysis,^2^ membrane transport,^3^ and ligand binding.^4^ In conventional molecular dynamics (cMD) simulations, however, protonation states are assigned before the simulation based on p*K*_a_ values estimated by empirical methods (e.g., PROPKA3^5^) or Poisson–Boltzmann methods (e.g., H++^6^ and DelPhiPKa^7^), and they are held fixed throughout the simulation. This approximation is valid only if the assigned protonation states are correct and do not change as the biomolecule samples conformations at the pH of interest. To address this limitation, constant-pH MD methods, which model protonation equilibria explicitly while propagating the atomic coordinates at a specified pH, have been developed over three decades (see ref 8 for a recent review).

Common constant-pH MD methods make use of either discrete or continuous representations of protonation states. In the discrete approach, also known as hybrid Monte Carlo (MC)/MD or stochastic constant-pH MD,^9^ conformational sampling is periodically interrupted by MC moves that attempt to switch protonation states. In this approach, the MC moves are evaluated with a continuum solvent model; it has been implemented^10,11^ for use with GROMACS^12^ and in Amber.^13,14^ A related approach^15^ avoids continuum electrostatics by switching protonation states gradually through alchemical transformations, which allows the solvent to relax and increases the acceptance rate of MC moves; this approach has been implemented^16^ in NAMD.^17^

This work focuses on the continuous constant-pH MD approach, hereafter referred to as CpHMD. In CpHMD, a set of titration coordinates (*λ* particles), whose end states correspond to distinct protonation states, is propagated simultaneously with the atomic coordinates using the *λ*-dynamics framework originally developed for free energy calculations.^18^ CpHMD was first developed in CHARMM for generalized Born (GB) implicit-solvent simulations (GB-CpHMD).^19,20^ A hybrid-solvent scheme^21^ was subsequently introduced to enable explicit-solvent conformational sampling while retaining the GB model for propagating titration particles. GB was later removed in CpHMD methods based on the generalized reaction field^22^ for long-range electrostatics or force switching.^23,24^ Finally, CpHMD methods that use particle-mesh Ewald (PME) electrostatics to propagate both the titration and conformational coordinates were developed in GROMACS,^25^ CHARMM,^26,27^ and Amber.^28^

The CpHMD approach can be implemented as a special case of *λ*-dynamics in which the potential energies of the end (protonation) states are interpolated. This approach is used in the CpHMD functionality of the multisite *λ*-dynamics (MS*λ*D) framework developed by Brooks and coworkers^23^ in CHARMM,^29^ and in an analogous implementation by Grubmüller and coworkers^25^ in GROMACS.^12^ The CpHMD approach can be alternatively implemented by interpolating the atomic charges of titratable sites, as in the original GB-CpHMD of Brooks and coworkers^19,20^ and the subsequent CpHMD implementations from our group.^21,22,26,28^ Charge interpolation is computationally more efficient and was recently adopted by Hess and coworkers^30^ to accelerate the PME-based CpHMD in GROMACS. Other differences among the three CpHMD implementations and their practical consequences will be discussed in future work. As major MD engines have been redesigned for GPU-based computing, recent CpHMD developments have focused on CUDA implementations, including GPU-accelerated MS*λ*D called BLaDE^27^ in CHARMM, PME-CpHMD_cuda_ ^28^ in Amber, and GPU-accelerated PME-CpHMD^30^ and fast multipole method (FMM)-CpHMD^31,32^ in GROMACS.

Here we report two coordinated developments in Amber: a split-kernel PME-CpHMD_cuda_ implementation that offers significant acceleration, and a native implementation of the asynchronous replica-exchange (AREX) algorithm^33^ that makes enhanced sampling accessible to laboratories with limited GPU resources. Additionally, we parameterized the model potential of mean force parameters for all six titratable residue types (Asp, Glu, His, Cys, Tyr, Lys; Tyr for the first time) for simulations with the CHARMM c22, Amber ff14sb and ff19sb force fields. We validated the new implementations and parameters with p*K*_a_ calculations of model peptides and benchmarked the performance on proteins ranging from 45 to 1,338 residues. The new PME-CpHMD_cuda_ incurs an overhead of 12–18% relative to cMD in the test cases, making it practical for routine studies of proton-coupled biomolecular processes. Finally, p*K*_a_ calculations of hen egg white lysozyme (HEWL) reveal force-field-dependent biases in the electrostatic description, highlighting remaining challenges.

## 2 Methods and Implementation

### 2.1 Split-kernel implementation of the nonbonded calculations

In our previous CUDA implementation^28^ of the all-atom PME-CpHMD,^26^ the CpHMD-specific changes to the direct-space electrostatic and Lennard-Jones (LJ) energies and forces were computed inside the pmemd.cuda nonbonded kernel. Because these changes involve only atoms in titratable groups, which make up a small fraction of the system, computing them in the nonbonded kernel adds overhead to the evaluation of atom pairs. To reduce the overhead, we moved the CpHMD-specific changes into a separate kernel, implemented in pmemd.cuda and pmemd.cuda.MPI in Amber24.^34^ We refer to the previous and new implementations as the fused- and split-kernel implementations, respectively.

Let {**r**_*i*_} denote the atomic coordinates. The nonbonded energy consists of the LJ, short-range direct-space Coulomb, and direct-space correction energies:^26^

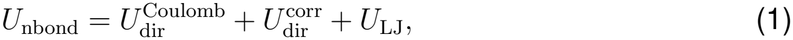

where

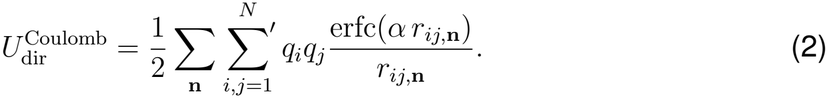

Here *N* is the number of atoms, *q*_*i*_ and *q*_*j*_ are the partial charges of atoms *i* and *j*, **n** runs over the periodic image lattice vectors, *r*_*ij*,**n**_ = |**r**_*i*_ − **r**_*j*_ + **n**| is the distance between atom *i* and the periodic image of atom *j* translated by **n**, *α* is the Ewald splitting parameter, and erfc is the complementary error function. The prime indicates that, for **n** = **0**, the terms with *i* = *j* and the terms for excluded pairs (*i, j*) *∈ M* are omitted. In practice, the sum is truncated at the real-space cutoff. For brevity, the Coulomb constant *κ* = 1*/*(4*πε*_0_) is omitted from all electrostatic energy expressions.

The direct-space correction energy consists of three terms,^26^

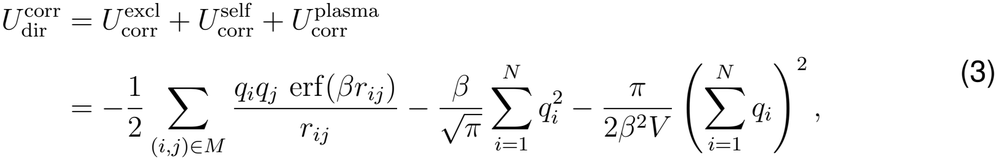

where *r*_*ij*_ = |**r**_*i*_ − **r**_*j*_|, erf is the error function, and *V* is the volume of the simulation box. The first term removes the interactions between excluded pairs included in the reciprocal-space sum. The second term removes the spurious interaction between a point charge and the compensating Gaussian charge included in the reciprocal-space sum. The third term corrects for the uniform background charge (plasma) that neutralizes a charged system.

In CpHMD, both 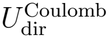 and 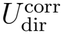 are *{λ*_*α*_}-dependent, because the partial charges of titratable groups are linearly interpolated between the protonated and deprotonated states:^19,20^

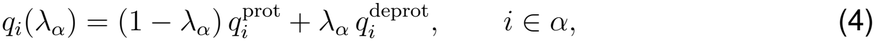

where index *α* denotes a titratable group. The LJ interactions involving titratable hydrogens are also *λ*-dependent:^19,20^

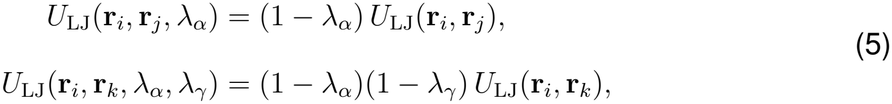

where the first equation describes the interaction between a titratable hydrogen *i* of group *α* and a nontitratable atom *j*, and the second describes the interaction between *i* and a titratable hydrogen *k* of a different group *γ*. The extensions of Eqs. 4 and 5 to tautomer titration are given in Ref.^20^

The nonbonded energy in CpHMD can therefore be written as

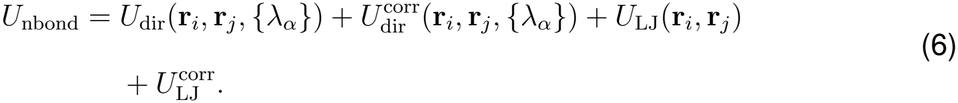

The first three terms are evaluated by the standard nonbonded kernel, whereas the CpHMD-specific LJ correction energy, defined as the difference between the *λ*-dependent and standard LJ energies of pairs involving titratable hydrogens, is computed in a new CpHMD kernel:

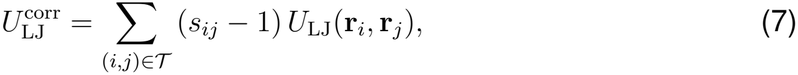

where *T* is the set of atom pairs involving at least one titratable hydrogen, and *s*_*ij*_ is (1*™λ*_*α*_) or (1 *™ λ*_*α*_)(1 *™ λ*_*γ*_), depending on whether one or both atoms are titratable hydrogens.

The derivative of the nonbonded energy with respect to the coordinates of atom *i* is

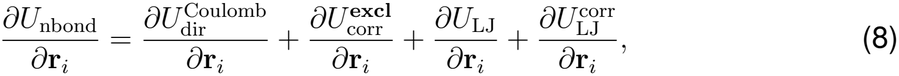

where the first three terms are evaluated by the standard PME nonbonded kernel, whereas the LJ force correction is evaluated by the new sparse CpHMD kernel.

The derivative of the nonbonded energy with respect to the titration coordinate *λ*_*α*_ is

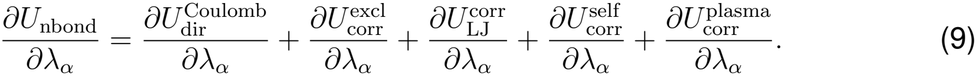

The first three terms are evaluated in the new sparse CpHMD kernel, whereas the last two terms are evaluated in the existing CpHMD self-correction kernel.

In practice, the direct-space atomic forces, energies, and, when required, the virial are first evaluated using the standard nonbonded kernel with *λ*-interpolated charges. A sparse CpHMD kernel then accumulates CpHMD-specific LJ corrections and atomic forces as well as the nonbonded contributions to the *λ* forces. To increase computational efficiency, neighbor-list entries containing atoms assigned to titratable groups are identified after each standard neighbor-list rebuild and stored in a compact list.

### 2.2 Native asynchronous pH replica exchange (AREX)

We implemented the asynchronous replica-exchange (AREX) algorithm^21,33^ in pmemd.cuda.MPI, separating a general exchange scheduler from the condition-specific exchange criterion.

Each replica is a copy of the system (coordinates, velocities, and *λ* values) that is assigned to one pH condition in the pH ladder; the scheduler can be generalized to other conditions such as temperature. The scheduler tracks the replica–condition assignments, the alternating exchange pattern, cycle completion, and checkpoint data, while the pH-REX module supplies the pH-dependent state variables and the Metropolis acceptance criterion.

At initialization, replicas are distributed among the available MPI process groups (workers). Each worker propagates its replica for one exchange cycle (typically 1000 MD steps) and returns the resulting state to the scheduler. As in the previous Python-based implementation,^33^ exchanges are attempted between neighboring conditions, alternating between the pairs (1, 2), (3, 4), … and (2, 3), (4, 5), … in successive exchange cycles. Unlike synchronous replica exchange, each exchange is evaluated as soon as both replicas in a pair have completed their segment, so idle workers can be reassigned without waiting for all replicas to finish.

Consider replicas *a* and *b*, currently at neighboring pH conditions pH_*a*_ and pH_*b*_. In the pH-REX method,^21^ an attempt to swap their pH conditions is accepted with probability

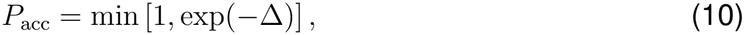

where Δ is the change in the total pH-biasing energy upon the swap,

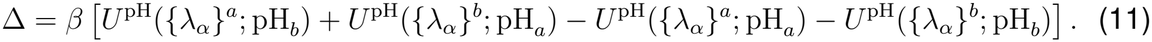

Here *β* = 1*/k*_*B*_*T*, and *{λ}*^*a*^ and *{λ}*^*b*^ are the titration coordinates of replicas *a* and *b*. The first two terms are the pH-biasing energies of the two replicas after the swap, and the last two are those before the swap. If the swap is accepted, replica *a* continues at pH_*b*_ and replica *b* at pH_*a*_; otherwise, both remain at their current pH. The pH-biasing energy is given as

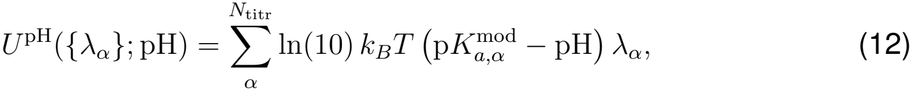

where *λ*_*α*_ and 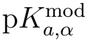 are the titration coordinate and model p*K*_*a*_ of residue *α*, respectively. *N*_titr_ is the total number of titratable residues.

This implementation transfers the full replica state (MD and titration data) explicitly between the scheduler and the workers. When a worker is assigned a replica, the replica’s state is loaded into the pmemd arrays and, where needed, the GPU; when the exchange cycle completes, the updated state is returned to the scheduler, which evaluates the exchange criterion.

Output files are tied to pH conditions. At initialization, a pH-labeled filename is derived from each pmemd output filename, and each pH condition keeps its own trajectory, CpHMD output, restart file, and trajectory time. At the start of each exchange cycle, the worker writes to the output files of its replica’s current pH condition. The exchange interval can be chosen independently of the trajectory and CpHMD output frequencies.

At the end of each exchange cycle, a replica can be continued in one of two modes. In continuous mode (the default), the worker resumes directly from the in-memory state, avoiding any perturbation from stopping and restarting. In restart-emulation mode (arex_emulate_resta selected transient MD and pH-REX quantities are reset, reproducing the previous Python-based implementation, in which each exchange interval was run as a separate pmemd.cuda job started from restart files.

Interrupted AREX simulations can be resumed from a global checkpoint file together with the per-condition restart files. The checkpoint file stores the scheduler state, as well as the output and pH-REX state of each replica. Each per-condition restart file stores the coordinates, velocities, box dimensions, and PHMD variables of the replica currently at that pH condition. Before the simulation resumes, the checkpoint is checked for consistency with the current system size, pH ladder, exchange criterion, and exchange interval. All AREX input options are listed in Table S1.

## 3 Simulation protocols

### 3.1 Model parameterization and validation

#### System preparation and TI simulations

Given the availability of NMR titration data,^35^ blocked tripeptides CH_3_CO-Gly-X-Gly-NH_2_ (X = Asp, Glu, Cys, His, Lys, or Tyr) were used as models. Simulation systems for use with the three force fields were built separately. First, simulation systems were built using CHARMM.^29^ A tripeptide was solvated in a truncated octahedral water box with at least 15 Å between the peptide and the box edge. One Na^+^ (Asp and Glu) or Cl^−^ (Lys) was added to neutralize charge at pH 7. The CHARMM files were converted to Amber format with chamber (AmberTools25)^34^ for simulations with the CHARMM (c22) force field. Next, equivalent systems were built in tleap for the Amber ff14sb and ff19sb force fields, with the number of water molecules matched to the c22 systems.

TI simulations of model peptides at fixed *θ* or *θ*_*x*_ values were performed to obtain the mean forces.^19,20^ After energy minimization, each system was heated from 100 to 300 K over 100 ps at constant volume, followed by equilibration at constant pressure, with harmonic restraints on heavy atoms reduced from 2.0 to 1.0 to 0.0 kcal mol^−1^ Å^−2^ in three 100 ps stages. The unrestrained production run lasted 20 ns, and the entire trajectory was used to calculate the mean force.

For the model peptides with single-site titration (Cys, Lys, and Tyr), mean forces at 15 *θ* values were obtained. For the model His peptide, mean forces were obtained at 16 (*θ, θ*_*x*_) values along each of three branches: varying *θ* at *θ*_*x*_ = 0 or *π/*2 and varying *θ*_*x*_ at *θ* = *π/*2. For model Asp and Glu peptides, a 9 *×* 14 (*θ, θ*_*x*_) grid was used. A total of 124 mean forces were obtained, with the remaining two supplied as analytical zero-force boundary conditions.

#### Force fields and MD protocol

Three force fields were used: c22/CMAP^36,37^ with CHARMM-modified TIP3P water,^36,38^ ff14sb^39^ with TIP3P water,^38^ and ff19sb^40^ with OPC water.^41^ For brevity, they are referred to as c22, ff14sb, and ff19sb, respectively. All simulations used a 2 fs time step (unless otherwise noted) with SHAKE constraints on bonds involving hydrogen,^42^ Langevin dynamics^43^ at 300 K with a collision frequency of 1 ps^−1^, and isotropic pressure coupling at 1 bar with a relaxation time of 0.1 ps. Consistent with the development of each force field, c22 simulations used a 12 Å nonbonded cutoff with force switching from 10 Å, whereas ff14sb and ff19sb simulations used cutoffs of 8 and 10 Å, respectively. Long-range electrostatics were treated with PME^44^ with a grid spacing of approximately 1 Å.

#### Parameter fitting

For model His peptide, the N*δ* and N*ϵ* titration, and the tautomer conversion profiles were fit simultaneously to the bivariate polynomial^20^ by a global fit with five parameters [*A*_1_, *C*_1_, *C*_0_, *A*_10_, *C*_10_], where *C*_1_ = *A*_1_*B*_1_, *C*_0_ = *A*_0_*B*_0_, and *C*_10_ = *A*_10_*B*_10_. These values were converted to *A*_0_, *B*_0_, *A*_1_, *B*_1_, *A*_10_, and *B*_10_ for the CpHMD parameter file.^20^

For model Asp and Glu peptides, a staged fit was used:^20^ *∂U/∂θ*_*x*_ was fit as a function of *θ*_*x*_ at each *θ*, and *∂U/∂θ* as a function of *θ* at each *θ*_*x*_. The resulting *A*(*θ*) values were fit to

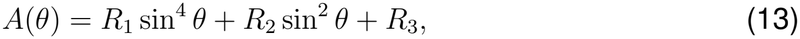

and *R*_4_ was fixed at 0.5 owing to the equivalence of the two carboxylate oxygens. For each fixed-*θ*_*x*_ slice, the coefficients of the *λ*^2^ and *λ* terms^20^ were calculated as

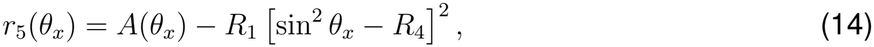

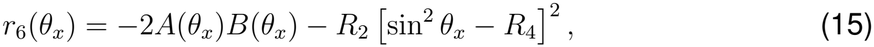

averaged over all slices, and converted to

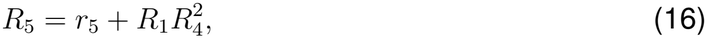

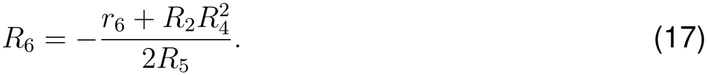

*R*_1_–*R*_6_ are included in the CpHMD parameter files.

#### Model peptide simulations and p*K*_a_ analysis

Three independent sets of AREX simulations were performed for each model peptide. Six pH values spaced by 0.5 units and bracketing the reference p*K*_a_ were used for each residue: Asp (2.5–5.0), Glu (3.0–5.5), His (5.5–8.0), Cys (7.5–10.0), Tyr (8.5–11.0), and Lys (9.0–11.5). Each pH replica was run for 21 ns; exchanges between adjacent replicas were attempted every 1000 steps (2 ps) in the last 20 ns. The initial 1 ns was excluded from analysis. Following our previous work,^28^ protonated and deprotonated states were defined as *λ ≤* 0.2 and *λ ≥* 0.8, respectively, and the deprotonated fraction was calculated as *S* = *N*_unprot_*/*(*N*_prot_ + *N*_unprot_). The *S* values were fit to the generalized Henderson–Hasselbalch (HH) equation to obtain the p*K*_a_ and Hill coefficient *n*.

### 3.2 Simulations of hen egg white lysozyme (HEWL)

#### System preparation and CpHMD protocol

The simulation system was prepared in CHARMM^29^ from chain A of PDB entry 2LZT, with an acetylated N-terminus and an amidated C-terminus. His residues were set to HSP, Asp and Glu were patched as ASPP2 and GLUP2,^20^ disulfide bonds were introduced with the DISU patch, and missing hydrogens were added with HBUILD. The protein was solvated in a truncated octahedral box with a minimum buffer of 15 Å (17,730 water molecules), and ions were added to neutralize the system (assuming standard protonation states) and reach an ionic strength of 150 mM (50 Na^+^ and 59 Cl^−^ ). The c22 simulation system with 50 mM ionic strength included 17 Na^+^, and 26 Cl^−^ . The system was converted to Amber format with chamber. The ff14sb and ff19sb systems were built with tleap, using the CpHMD-specific AS2 and GL2 templates for Asp and Glu, HIP for His, and CYX for disulfide-bonded Cys, with the same numbers of water molecules and ions as the c22 system.

Each system was minimized for 10,000 steps (2000 steepest descent and 8000 conjugate gradient) and then heated from 100 to 300 K over 1 ns at constant volume, using a 1 fs time step with CpHMD at pH 7.5. Protein heavy atoms were restrained with a force constant of 100 kcal mol^−1^ Å^−2^. Each system was equilibrated for 3 ns at 18 pH values (0.5–9.0 in increments of 0.5) while the restraint force constant was reduced step-wise: 100, 10, 1, 0.1, and 0 kcal mol^−1^ Å^−2^ during 0–0.1, 0.1–0.2, 0.2–1.5, 1.5–2.0, and 2.0–3.0 ns, respectively. Unrestrained AREX production was then run for 150 ns per pH replica, with all other settings as in the model-peptide simulations.

#### Finite-size p*K*_a_ corrections

In PME simulations, the average electrostatic potential of the simulation box is set to zero, which leaves out the potential that arises from the trace of the solvent quadrupole moment, also known as the Bethe potential:^45–47^

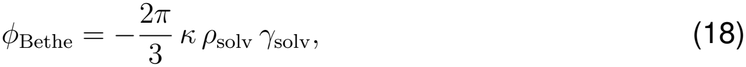

where *ρ*_solv_ = *N*_solv_*/⟨V*_box_*⟩* is the solvent number density in the simulation box, *γ*_solv_ is the quadrupole-moment trace of a solvent molecule, and *κ* is the Coulomb constant. Since *ρ*_solv_ is defined with respect to the total box volume, which includes the volume occupied by the solute, the missing potential depends on the box size. This was introduced as the type-D finite-size correction for PME-based free energy calculations.^48^

In PME-CpHMD simulations, the p*K*_a_ value is calculated relative to the model peptide in solution. If solvent number density differs between the protein and model-peptide systems, which is often the case, a correction is needed:^26,28^

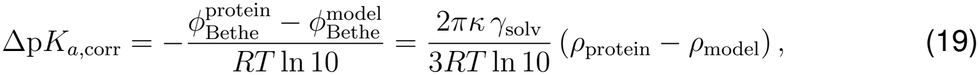

where *R* is the gas constant, *T* is the simulation temperature, and *κ* = 332.06kcalmol^−1^*e*^−2^. With the oxygen atom as the origin, *γ*_solv_ is calculated as 0.764 and 0.999 *e* Å^2^ for the TIP3P^38^ and OPC^41^ water models, respectively. Applying Equation 19, the p*K*_a_ corrections for Asp, Glu, and His residues in HEWL were −0.40 units with c22; −0.37 with ff14sb; and −0.54 with ff19sb (Table S7).

#### Coupled titration of E35/D52

A coupled titration pair has four microstates: unprotonated (E35^−^/D52^−^), singly protonated (E35H/D52^−^ or E35^−^/D52H), and doubly protonated (E35H/D52H). The average number of protons bound to the pair can be described by a two-step protonation model,^3,49^

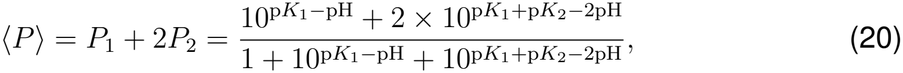

where *P*_1_ and *P*_2_ are the probabilities of the singly and doubly protonated states, and p*K*_1_ and p*K*_2_ are the macroscopic p*K*_a_’s for binding the first and second protons, respectively (p*K*_1_ *≥* p*K*_2_). The observed counts of the zero-, one-, and two-proton states across all pH conditions were fitted simultaneously to the two-step protonation model by multinomial maximum likelihood, with the constraint p*K*_1_ *≥* p*K*_2_.

## 4 Results and Discussion

### 4.1 Performance of the split-kernel PME-CpHMD

We benchmarked the throughput of the split-kernel PME-CpHMD implementation against the previous fused-kernel implementation and cMD using four protein systems, spanning approximately 30,000 to 281,000 atoms, namely, the soluble proteins BBL and HEWL and transmembrane proteins, a lipid transporter TacF and an ion channel ASIC trimer. With a 2 fs time step, the split-kernel PME-CpHMD achieved from 96 ns/day for the ASIC trimer to 378 ns/day for BBL. Hydrogen mass repartitioning (HMR)^50,51^ with a 4 fs time step increased the throughput by 75–82%, to 174–660 ns/day (Table 1 and Figure 1a). Relative to cMD, the split-kernel implementation reached 82–88% of the throughput, compared with 42–59% for the fused-kernel implementation (Figure 1b and Table 1), and exceeded the 60–70% reported for the recent GPU implementation of PME-CpHMD in GROMACS.^30^

**Table 1:**
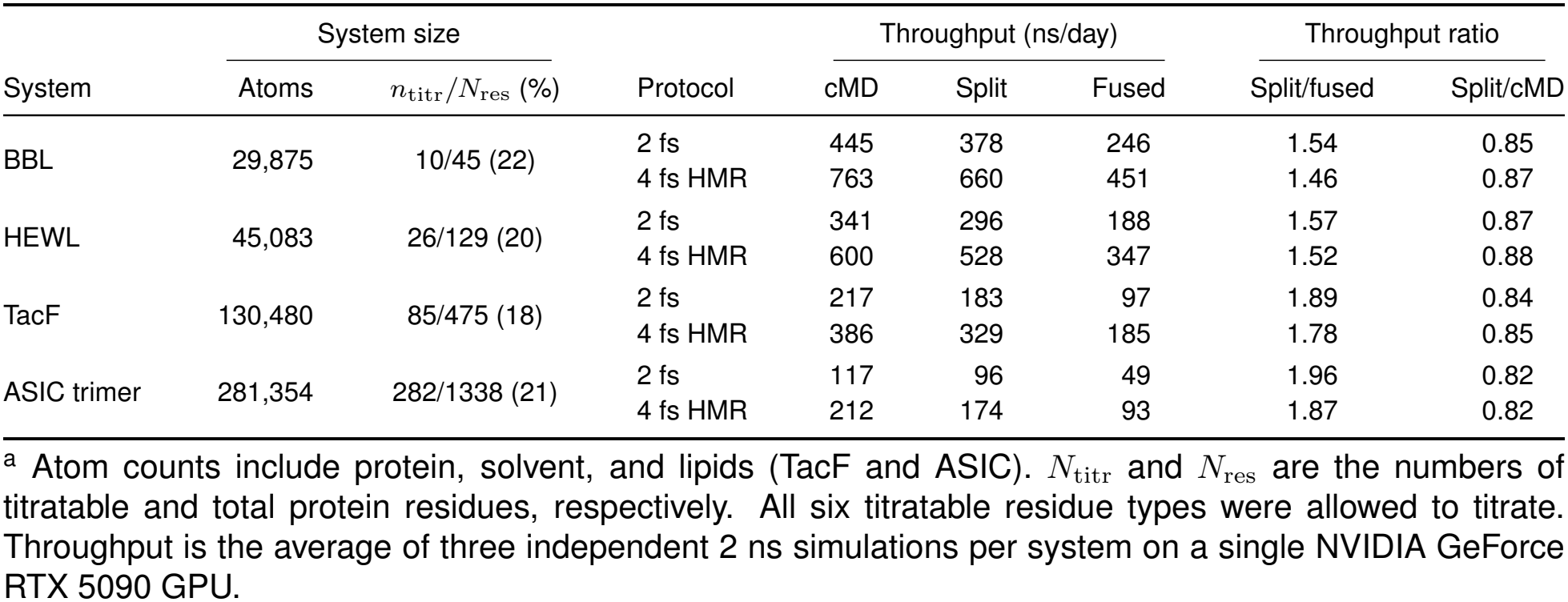
Single-GPU performance of the new split-kernel PME-CpHMD compared to the previous fused-kernel implementation and cMD^a^.

|  | System size |  |  | Throughput (ns/day) |  |  | Throughput ratio |  |
|---|---|---|---|---|---|---|---|---|
| System | Atoms | $n_{\mathrm{titr}}/N_{\mathrm{res}}$ (%) | Protocol | cMD | Split | Fused | Split/fused | Split/cMD |
| BBL | 29,875 | 10/45 (22) | 2 fs | 445 | 378 | 246 | 1.54 | 0.85 |
|  |  |  | 4 fs HMR | 763 | 660 | 451 | 1.46 | 0.87 |
| HEWL | 45,083 | 26/129 (20) | 2 fs | 341 | 296 | 188 | 1.57 | 0.87 |
|  |  |  | 4 fs HMR | 600 | 528 | 347 | 1.52 | 0.88 |
| TacF | 130,480 | 85/475 (18) | 2 fs | 217 | 183 | 97 | 1.89 | 0.84 |
|  |  |  | 4 fs HMR | 386 | 329 | 185 | 1.78 | 0.85 |
| ASIC trimer | 281,354 | 282/1338 (21) | 2 fs | 117 | 96 | 49 | 1.96 | 0.82 |
|  |  |  | 4 fs HMR | 212 | 174 | 93 | 1.87 | 0.82 |

<sup>a</sup> Atom counts include protein, solvent, and lipids (TacF and ASIC). $N_{\mathrm{titr}}$ and $N_{\mathrm{res}}$ are the numbers of titratable and total protein residues, respectively. All six titratable residue types were allowed to titrate. Throughput is the average of three independent 2 ns simulations per system on a single NVIDIA GeForce RTX 5090 GPU.

**Table 2:**
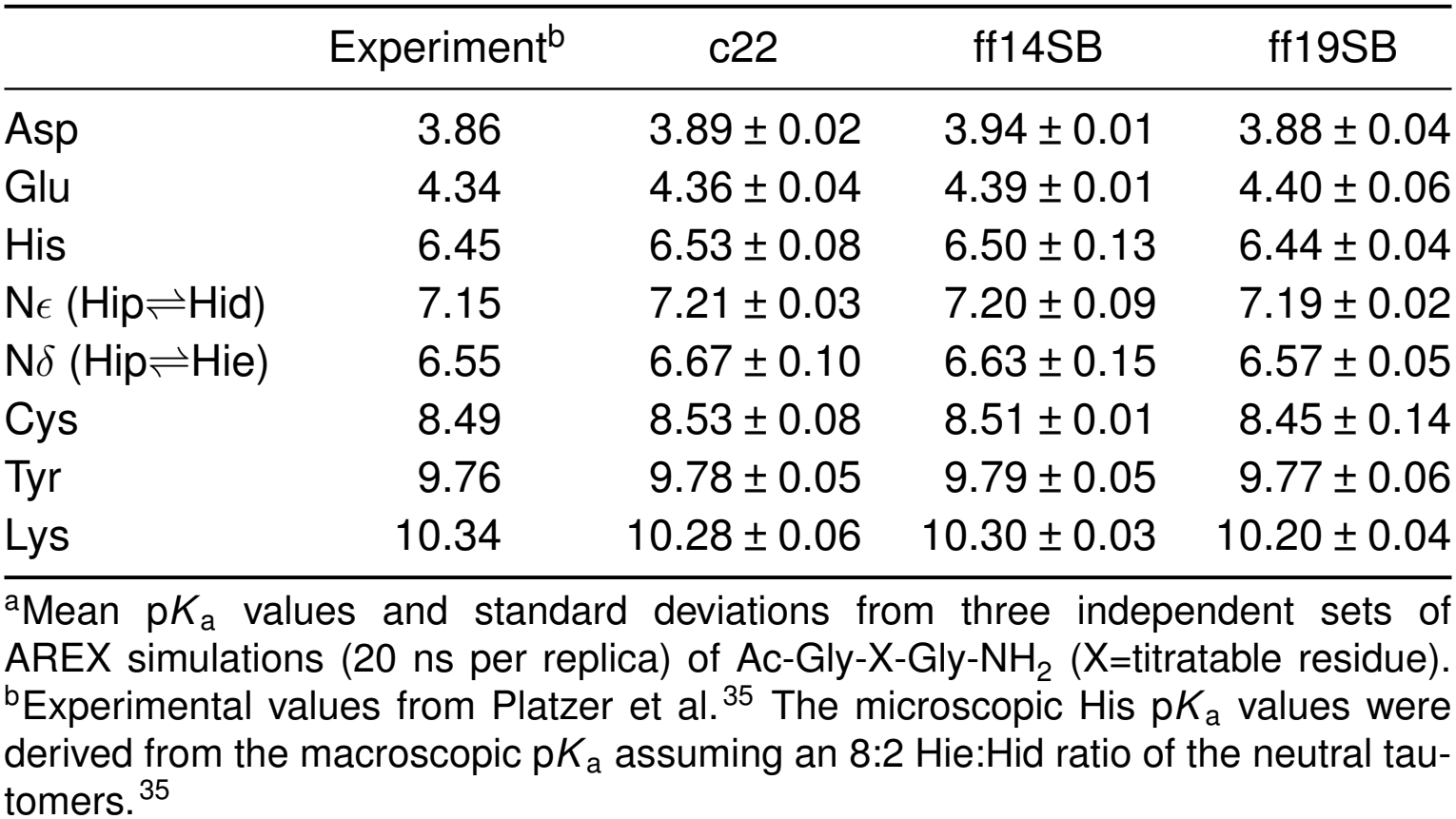
Calculated and experimental p*K*_a_ values of model tripeptides^a^.

**Figure 1:**
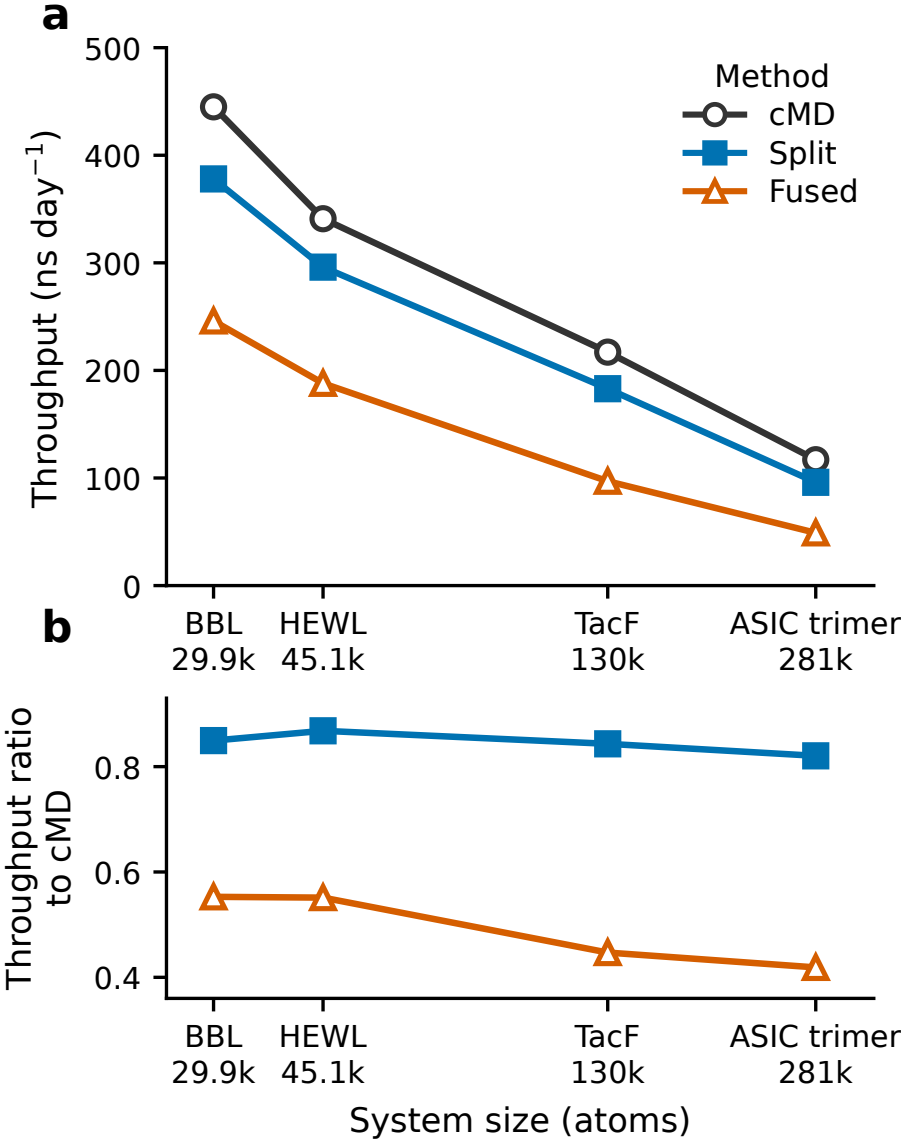
Single-GPU performance of split-kernel PME-CpHMD compared with fused-kernel implementation and cMD. **(a)** Throughput of cMD (black), split-kernel (blue), and fused-kernel (orange) PME-CpHMD. **(b)** Throughput relative to cMD. All simulations used a 2-fs time step and identical simulation settings on a single NVIDIA GeForce RTX 5090 GPU. Throughput was calculated from the wall-clock times of three independent 2 ns production runs per system, with relative standard deviations below 0.8%. Numerical values and additional comparisons are given in Table 1.

At these throughputs, a 1 µs PME-CpHMD simulation with HMR on a single RTX 5090 would take approximately 1.5 days for BBL, 1.9 days for HEWL, 3.0 days for TacF, and 5.7 days for the ASIC trimer, making microsecond-scale PME-CpHMD simulations routinely feasible on a single consumer-grade GPU.

### 4.2 Performance of the native AREX implementation

To verify that results are independent of the number of GPUs, we performed ten sets of AREX titration simulations of the Asp tripeptide with one, two, three, and four GPUs. The mean calculated p*K*_a_ values were 3.90, 3.91, 3.90, and 3.90, respectively, with standard deviations of 0.04–0.06 (Figure 2a and Table S5). Sampling was likewise unaffected: the exchange acceptance ratio was 80.9–81.1%, the mean number of round trips per replica across the pH ladder was 252.5–263.4, and all 240 replicas visited every pH condition. Thus, the number of GPUs does not affect the calculated p*K*_a_ or the sampling of the pH ladder.

**Figure 2:**
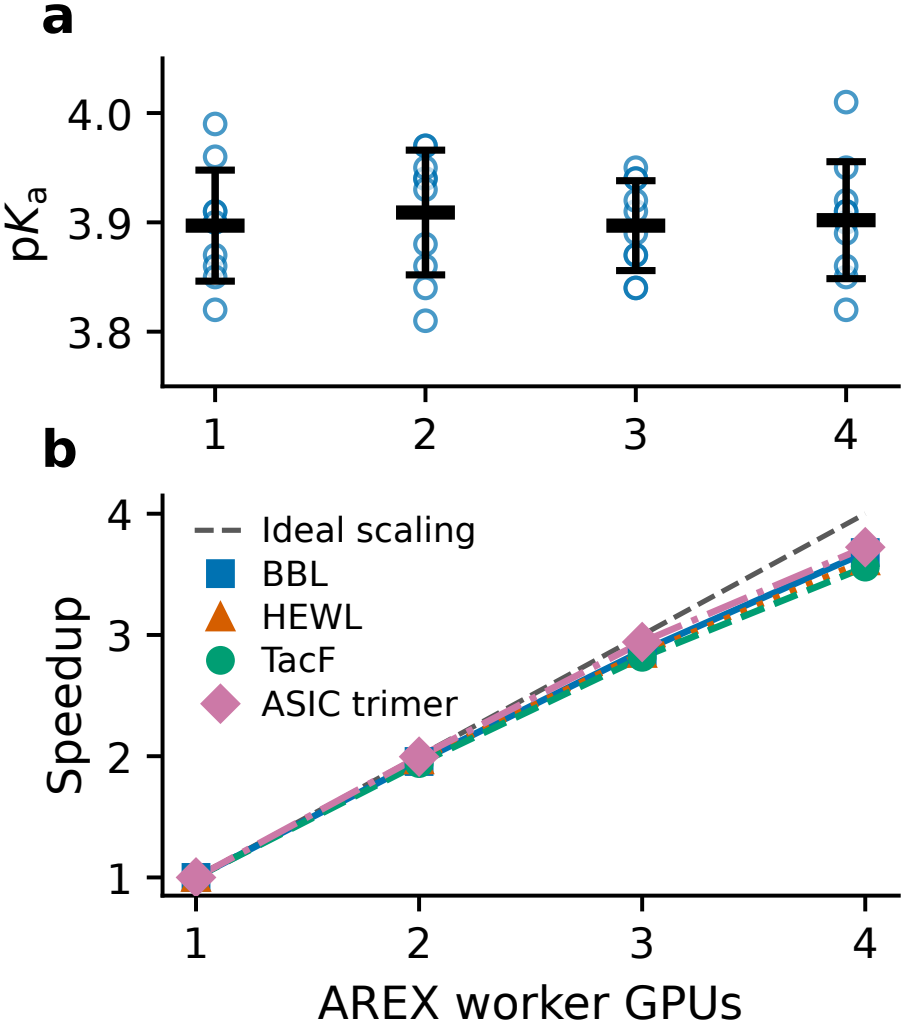
GPU-count validation and multi-GPU scaling of AREX. **(a)** Calculated p*K*_a_ of the Asp tripeptide from ten sets of AREX simulations with one to four GPUs. Open circles show individual simulations, horizontal bars show means, and error bars show standard deviations. Each simulation was run for 21 ns per replica, and the first 1 ns was discarded. **(b)** Speedup of AREX relative to one GPU for BBL, HEWL, TacF, and the ASIC trimer. Each simulation used 24 pH replicas (pH 0.5–12.0 in increments of 0.5) and was run for 2 ns per replica, titrating all Asp, Glu, His, Cys, Tyr, and Lys. The dashed line indicates ideal linear scaling.

We also evaluated the multi-GPU scaling of AREX on one to four GPUs for BBL, HEWL, TacF, and ASIC trimer. With two, three, and four GPUs, the parallel efficiencies were 97.0–99.8%, 93.9–98.1%, and 89.0–93.1%, respectively, corresponding to four-GPU speedups of 3.56–3.72 (Table S6). The modest decrease in efficiency with increasing GPU count is likely due to the growing overhead of inter-GPU communication.

### 4.3 Parameterization and validation of model p*K*_a_’s

Leveraging the speedup of the new PME-CpHMD implementation, we determined the model titration parameters for Asp, Glu, His, Cys, and Lys and derived new parameters for Tyr for the c22, ff14sb, and ff19sb force fields. Blocked tripeptides, Ac-Gly-X-Gly-NH_2_ (X=titratable residue), were used as models, with reference p*K*_a_ values taken from Platzer et al.^35^ Compared with the previous protocol,^28^ we increased the number of *λ* windows for TI simulations and revised the fitting procedure (Simulation protocols and Supp. Table S1–S5).

Across the six tripeptides and three force fields (24 p*K*_a_ values in total, including the microscopic p*K*_a_’s for the two His tautomers), the mean calculated p*K*_a_ values deviated from experiment by at most 0.14 units. The new Tyr parameters gave absolute deviations of 0.02, 0.09, and 0.08 units for c22, ff14sb, and ff19sb, respectively. The largest deviation was for Lys with ff19sb, for which the calculated p*K*_a_ is 0.14 units lower than the experimental value. The largest standard deviation among replicates (0.15 units) was for the microscopic p*K*_a_ of Hid (Hip *⇀↽* Hie) with ff14sb. Note that no adjustment of parameters was made. These results confirm that the new model titration potentials of mean force consistently recover the experimental model p*K*_a_ values. The p*K*_a_ values and Hill coefficients for each replicate, the cumulative deprotonated fractions for replicate 1, and the titration curves for all three replicates are provided in the Supporting Information.

### 4.4 p*K*_a_ calculation for HEWL with CHARMM and Amber force fields

#### Overall assessment

To further validate the split-kernel PME-CpHMD implementation and to examine the effect of the force field, we calculated the p*K*_a_ values of hen egg white lysozyme (HEWL) from AREX simulations with the c22, ff14sb, and ff19sb force fields. Each simulation was run for 150 ns per replica, and the last 50 ns were used for p*K*_a_ calculations and analysis (Table 3 and Suppl. Figure S6–S11). The root-mean-square deviations (RMSDs) from experiment^52^ are 0.93, 1.26, and 1.15 for c22, ff14sb, and ff19sb, respectively, with corresponding Pearson correlation coefficients (*r*) of 0.88, 0.73, and 0.69.

**Table 3:**
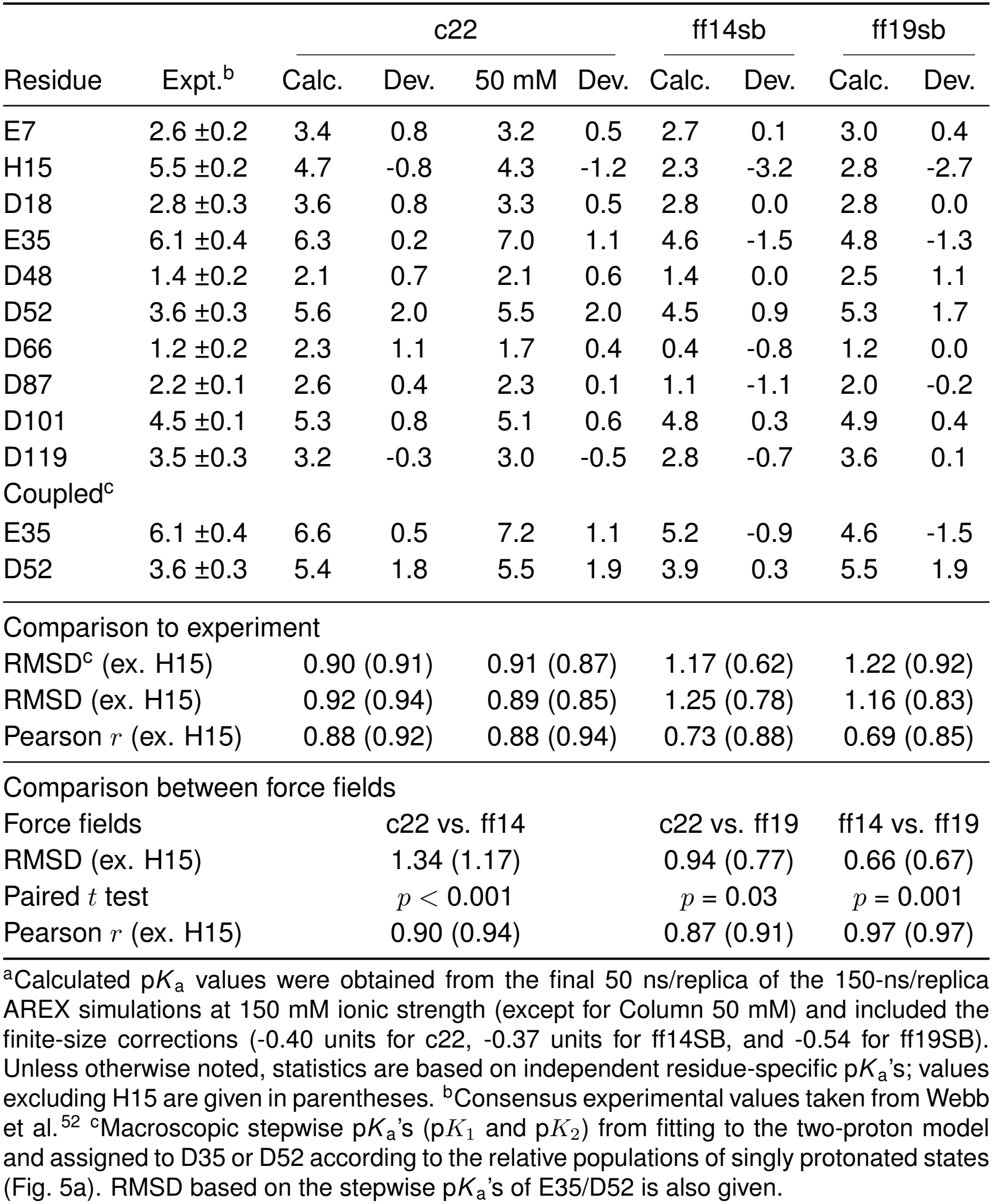
Experimental and calculated p*K*_a_ values of hen egg white lysozyme (HEWL) using three force fields.^a^.

|  |  | c22 |  |  |  | ff14sb |  | ff19sb |  |
|---|---|---|---|---|---|---|---|---|---|
| Residue | Expt.<sup>b</sup> | Calc. | Dev. | 50 mM | Dev. | Calc. | Dev. | Calc. | Dev. |
| E7 | 2.6 ±0.2 | 3.4 | 0.8 | 3.2 | 0.5 | 2.7 | 0.1 | 3.0 | 0.4 |
| H15 | 5.5 ±0.2 | 4.7 | -0.8 | 4.3 | -1.2 | 2.3 | -3.2 | 2.8 | -2.7 |
| D18 | 2.8 ±0.3 | 3.6 | 0.8 | 3.3 | 0.5 | 2.8 | 0.0 | 2.8 | 0.0 |
| E35 | 6.1 ±0.4 | 6.3 | 0.2 | 7.0 | 1.1 | 4.6 | -1.5 | 4.8 | -1.3 |
| D48 | 1.4 ±0.2 | 2.1 | 0.7 | 2.1 | 0.6 | 1.4 | 0.0 | 2.5 | 1.1 |
| D52 | 3.6 ±0.3 | 5.6 | 2.0 | 5.5 | 2.0 | 4.5 | 0.9 | 5.3 | 1.7 |
| D66 | 1.2 ±0.2 | 2.3 | 1.1 | 1.7 | 0.4 | 0.4 | -0.8 | 1.2 | 0.0 |
| D87 | 2.2 ±0.1 | 2.6 | 0.4 | 2.3 | 0.1 | 1.1 | -1.1 | 2.0 | -0.2 |
| D101 | 4.5 ±0.1 | 5.3 | 0.8 | 5.1 | 0.6 | 4.8 | 0.3 | 4.9 | 0.4 |
| D119 | 3.5 ±0.3 | 3.2 | -0.3 | 3.0 | -0.5 | 2.8 | -0.7 | 3.6 | 0.1 |
| Coupled<sup>c</sup> |  |  |  |  |  |  |  |  |  |
| E35 | 6.1 ±0.4 | 6.6 | 0.5 | 7.2 | 1.1 | 5.2 | -0.9 | 4.6 | -1.5 |
| D52 | 3.6 ±0.3 | 5.4 | 1.8 | 5.5 | 1.9 | 3.9 | 0.3 | 5.5 | 1.9 |
| Comparison to experiment |  |  |  |  |  |  |  |  |  |
| RMSD<sup>c</sup> (ex. H15) |  | 0.90 (0.91) |  | 0.91 (0.87) |  | 1.17 (0.62) |  | 1.22 (0.92) |  |
| RMSD (ex. H15) |  | 0.92 (0.94) |  | 0.89 (0.85) |  | 1.25 (0.78) |  | 1.16 (0.83) |  |
| Pearson *r* (ex. H15) |  | 0.88 (0.92) |  | 0.88 (0.94) |  | 0.73 (0.88) |  | 0.69 (0.85) |  |
| Comparison between force fields |  |  |  |  |  |  |  |  |  |
| Force fields |  | c22 vs. ff14 |  |  |  | c22 vs. ff19 |  | ff14 vs. ff19 |  |
| RMSD (ex. H15) |  | 1.34 (1.17) |  |  |  | 0.94 (0.77) |  | 0.66 (0.67) |  |
| Paired *t* test |  | *p* < 0.001 |  |  |  | *p* = 0.03 |  | *p* = 0.001 |  |
| Pearson *r* (ex. H15) |  | 0.90 (0.94) |  |  |  | 0.87 (0.91) |  | 0.97 (0.97) |  |

<sup>a</sup>Calculated p*K*<sub>a</sub> values were obtained from the final 50 ns/replica of the 150-ns/replica AREX simulations at 150 mM ionic strength (except for Column 50 mM) and included the finite-size corrections (-0.40 units for c22, -0.37 units for ff14SB, and -0.54 for ff19SB). Unless otherwise noted, statistics are based on independent residue-specific p*K*<sub>a</sub>'s; values excluding H15 are given in parentheses. <sup>b</sup>Consensus experimental values taken from Webb et al.<sup>52</sup> <sup>c</sup>Macroscopic stepwise p*K*<sub>a</sub>'s (p*K*<sub>1</sub> and p*K*<sub>2</sub>) from fitting to the two-proton model and assigned to D35 or D52 according to the relative populations of singly protonated states (Fig. 5a). RMSD based on the stepwise p*K*<sub>a</sub>'s of E35/D52 is also given.

We next examined how much the calculated p*K*_a_ values differ between force fields (Table 3). As expected, the two Amber force fields agree most closely, with a root-mean-square deviation (RMSD) of 0.66 and *r* = 0.97. By comparison, the RMSDs between c22 and ff14sb and between c22 and ff19sb are 1.34 and 0.94, respectively, with *r* values of 0.90 and 0.87, suggesting that the c22 results are closer to those of ff19sb than to those of ff14sb. Excluding H15 (discussed below) lowers the RMSDs between c22 and ff14sb and between c22 and ff19sb to 1.17 and 0.77, respectively, and raises the corresponding *r* values to 0.94 and 0.91, but the trend remains unchanged. Below we focus the discussion on H15, which shows the largest deviations from experiment, and on the E35/D52 catalytic dyad, which exhibits coupled titration.

#### p*K*_a_ calculation of the buried H15

All three force fields underestimate the p*K*_a_ of the partially buried H15 (Figure 3a), but the magnitude of error is much larger with ff14sb and ff19sb (3.2 and 2.7 units, respectively) than with c22 (0.8 units). Both the underes-timation and the force-field trend are consistent with the calculated p*K*_a_’s of the partially buried H166 in BBL using these three force fields.^53^ To characterize the degree of solvent sequestration, we calculated the fractional solvent-accessible surface area (*f*_SASA_) of the H15 side chain relative to that in solution, obtained from model tripeptide simulations. As for H166 in BBL,^53^ the pH profiles of the buried fraction (1 − fSASA) and the deprotonated fraction of H15 are strongly correlated (Figure 3b,c): as H15 becomes protonated and charged, its solvent exposure increases and its buried fraction decreases. In the simulations with the Amber force fields, both protonation and the increase in solvent exposure begin at lower pH. Importantly, while the buried fraction is similar between c22 and the Amber force fields for neutral H15, it differs for the charged form: as H15 becomes protonated, it remains consistently more buried in the ff14sb and ff19sb simulations than in the c22 simulations (Figure 3c). This suggests that the Amber force fields may underestimate the solvation of charged histidine relative to c22, disfavoring the protonated state and thereby exaggerating the p*K*_a_ downshift due to the desolvation penalty.

**Figure 3:**
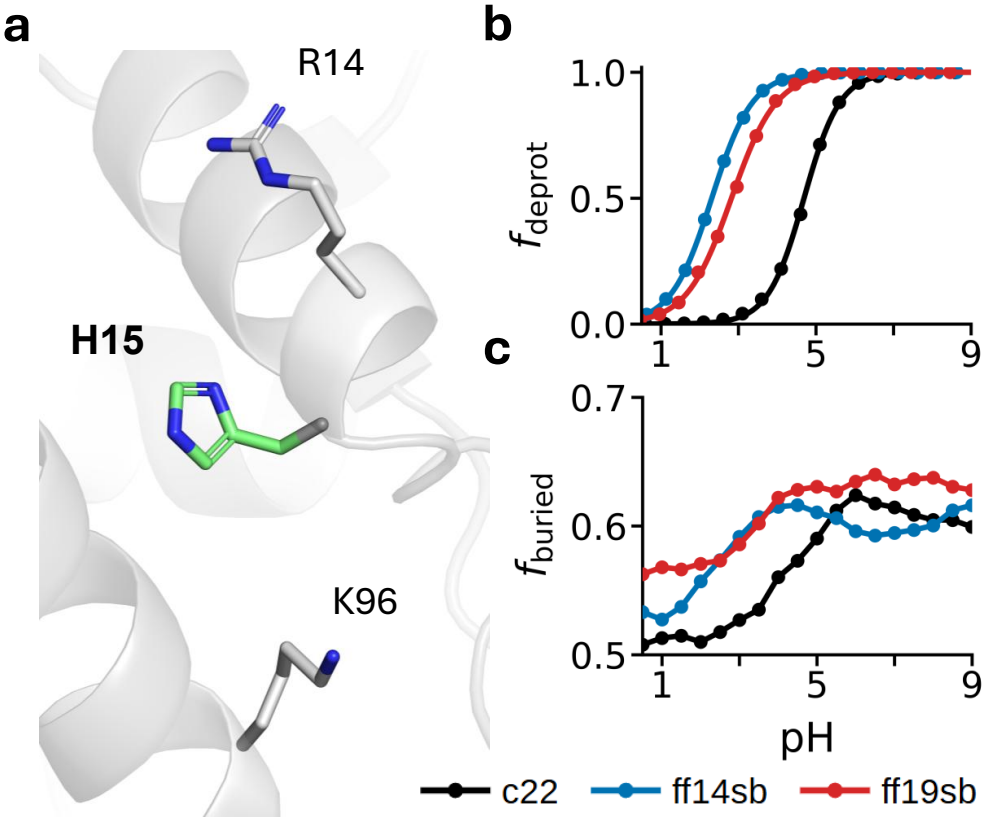
Titration and solvent exposure of H15 in HEWL from the c22, ff14sb, and ff19sb simulations. **a.** A snapshot of HEWL highlighting the H15 side chain (green) and nearby residues R14 and K96. **b**. Unprotonated fraction of H15 as a function of pH from the c22 (black), ff14sb (blue), and ff19sb (red) simulations. Symbols represent simulation data, and solid curves are best fits to the generalized HH equation. **c**. The buried fraction of H15 (1 − fSASA) as a function of pH. fSASA was calculated as the solvent accessible surface area (SASA) of the H15 sidechain relative to the corresponding model-peptide in solution.

#### Independent titration of E35/D52 in HEWL

We next examined the catalytic dyad E35/D52 (Figure 4a), which have divergent p*K*_a_’s between force fields. When treated independently, E35/D52 have p*K*_a_values of 6.3/5.6 (c22), 4.6/4.5 (ff14sb), 4.8/5.3 (ff19sb), respectively (Table 3). The distinct force-field dependence is further demonstrated by the independent E35 and D52 titration curves (Figure 4b). E35 deprotonates at substantially lower pH with both Amber force fields compared to c22. In contrast, for D52 this trend is only seen with ff14sb, while ff19sb is largely in agreement with c22.

**Figure 4:**
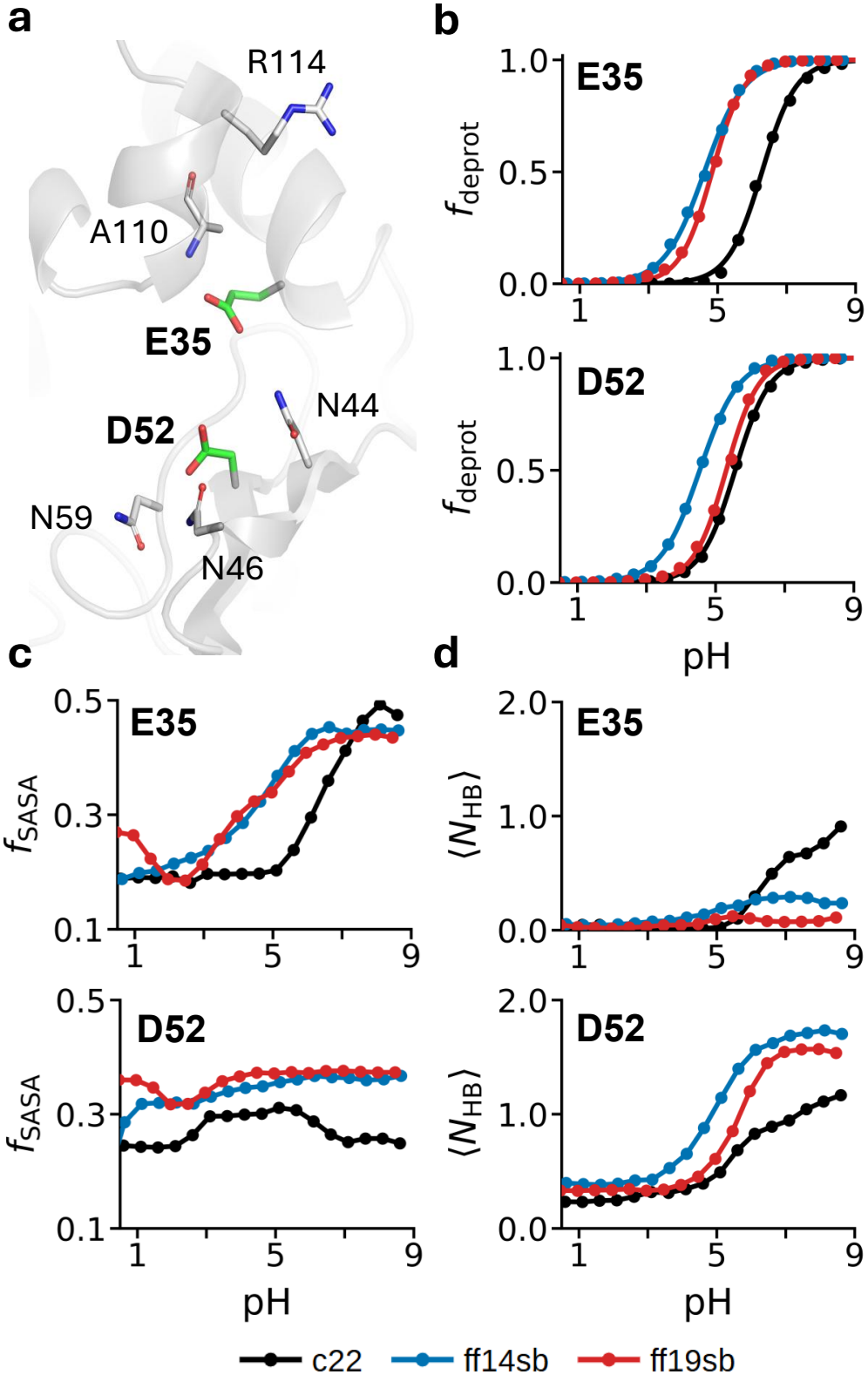
Titration and electrostatic environments of E35 and D52 in HEWL from the PME-CpHMD simulations with the c22, ff14sb, and ff19sb force fields. **a.** Snapshot of the active site of HEWL from a representative simulation frame, showing E35 and D52 (green) and (potential) h-bonding partners A110, R114, N44, N46, and N59. **b**. pH-dependent deprotonated fraction (*f*_deprot_) of E35 (top) and D52 (bottom) from the c22 (black), ff14sb (blue), and ff19sb (red) simulations. Symbols are simulation data, and curves are best fits to the generalized HH equation. **c**. pH-dependent fractional solvent exposed surface area (*f*_SASA_) of the carboxylate oxygens relative to the value of the corresponding model tripeptide in solution. **d**. The number of h-bonds (*N*_HB_) accepted by E35 (A110 and R114) and D52 (N44, N46, and N59) as a function of pH.

To understand the force-field effects, we examined the pH dependence of *f*_SASA_ and the number of hydrogen bonds (h-bonds, *N*_HB_), and compared them with the pH dependence of the deprotonated fraction (*f*_deprot_; Figure 4b–d; Suppl. Figure S14). For E35, *f*_SASA_ covers a similar range with all three force fields. However, as with *f*_deprot_, the *f*_SASA_ profile in the c22 simulation is shifted to a higher pH range. Importantly, within the titration range (between the protonated and deprotonated states), E35 is less solvent-exposed in the c22 simulation than in the ff14sb and ff19sb simulations. E35 is not involved in significant h-bonding in simulations with either Amber force field (Figure 4d). In the c22 simulation, deprotonated E35 forms an h-bond with R114; however, this only occurs outside of the titration range (pH *>* 6.4). These data suggest that the higher p*K*_a_ of E35 from the c22 simulation can be mainly attributed to its reduced solvent exposure.

For D52, the pH-dependent *N*_HB_ profile follows that of *f*_deprot_ with all three force fields, whereas *f*_SASA_ shows little pH dependence. With ff14sb, both *N*_HB_ and *f*_deprot_ profiles are shifted toward lower pH, and D52 forms more h-bonds within the titration range than with ff19sb and c22. The solvent exposure of D52 is similar with the two Amber force fields and slightly lower with c22. These results suggest that the lower p*K*_a_ of D52 with ff14sb compared to c22 and ff19sb may be attributed to the enhanced h-bond formation, with D52 as the acceptor, which stabilizes its deprotonated form.

#### Coupled titration of E35/D52 in HEWL

Since the minimum distance between the carboxylate groups of E35 and D52 is about 6.1 Å in the X-ray crystal structure (PDB entry 2LZT), the side chains may be electrostatically coupled (Suppl. Figure S12–S13). We therefore fitted the total number of protons bound to the dyad to the two-step protonation model (Eq. 20). The resulting stepwise p*K*_a_ values p*K*_1_/p*K*_2_ are 6.6/5.4, 5.2/3.9, and 5.5/4.6 for the c22, ff14sb, and ff19sb simulations, respectively. For both c22 and ff19sb, the stepwise p*K*_a_ values are about 0.2–0.3 units different from the independent residue-specific p*K*_a_ values, indicating that the coupling is weak. In contrast, electrostatic coupling is strongest in the ff14sb simulation: while the independent p*K*_a_ values differ by only 0.1 units, p*K*_1_ and p*K*_2_ differ by 1.3 units.

To assign the stepwise p*K*_a_’s to individual residues, we analyzed the pH-dependent fractions of the four joint microscopic protonation states of E35 and D52: E35^−^/D52^−^, E35H/D52^−^, E35^−^/D52H, and E35H/D52H (Figure 5a). The stepwise p*K*_a_’s can be assigned by comparing the fractions of the two singly protonated states, E35H/D52^−^ and E35^−^/D52H. For c22, E35H/D52^−^ dominates, accounting for 84% of the singly protonated population, which indicates that E35 has the higher p*K*_a_. This agrees with the independent p*K*_a_ calculation and with experiment, which identifies E35 as the proton donor of the catalytic dyad.^52^ For ff14sb, E35H/D52^−^ is only slightly more populated than E35^−^/D52H (by 6 %), consistent with the independent p*K*_a_ of E35 being only 0.1 unit higher than that of D52. Nonetheless, because E35H/D52^−^ is the preferred state, the higher stepwise value, p*K*_1_, can be assigned to E35. Surprisingly, with ff19sb, the protonation preference is reversed: only 29% of the singly protonated population is E35H/D52^−^. Therefore, D52 is assigned p*K*_1_, which is consistent with the independent p*K*_a_ calculation but contradicts experiment. Together, these results show that both the strength of the electrostatic coupling and the microscopic composition of the singly protonated population are highly sensitive to the force field.

**Figure 5:**
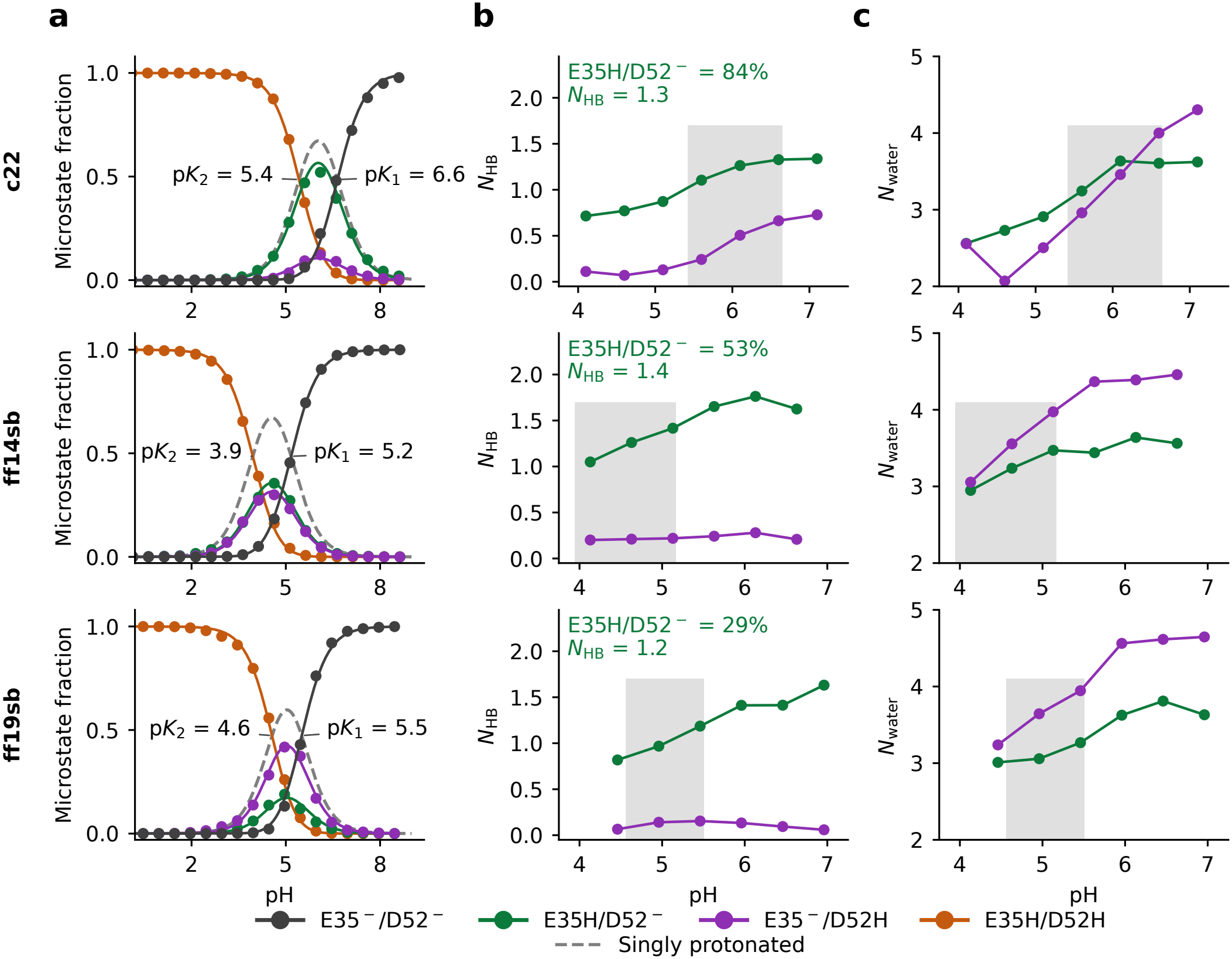
Coupled titration and microstate-conditioned environments of the catalytic dyad E35/D52 in HEWL from the PME-CpHMD simulations with the c22, ff14sb, and ff19sb force fields. **a.** pH-dependent fraction of the four microstate populations, E35H/D52H, E35H/D52^−^, E35^−^/D52H, and E35^−^/D52^−^. Curves are best fits to the four-state model. The dashed grey curve is the total population of the two singly protonated states; its crossings with E35^−^/D52^−^ and E35H/D52H give p*K*_1_ and p*K*_2_, respectively, obtained from fitting to the two-proton model (Eq. 20). **b**. Number of h-bonds (*N*_HB_) accepted by the deprotonated carboxylate in E35^−^/D52H (purple) or E35H/D52^−^ (green). The percentage indicates the fraction of E35H/D52^−^ within the singly protonated population. The annotated *N*_HB_ is the mean number of h-bonds for E35H/D52^−^ (green) at p*K*_1_. **c**. Number of water molecules (*N*_water_) within 3.4 Å of the deprotonated carboxylate oxygens (first-shell) in each singly protonated state. In panels **b** and **c**, the shaded region spans p*K*_2_ to p*K*_1_. All pH values include the force-field-specific p*K*_a_ corrections. H-bond is defined by a donor–acceptor distance *≤* 3.5 Å and an angle *≥* 150^*°*^.

To determine why each force field favors a particular singly protonated microstate, we analyzed the number of h-bonds and the number of first solvation shell water molecules in each of the two microstates (Figure 5b–c). Consistent with the residue-specific analysis (Figure 4), h-bonding favors E35H/D52^−^ over E35^−^/D52H with all three force fields (Figure 5b). In terms of solvent accessibility, c22 shows negligible difference between the two microstates or a slight preference of E35H/D52^−^, whereas both Amber force fields show a modest preference for E35^−^/D52H (Figure 5c). Thus, the microstate preference is determined by h-bonding alone with c22, corroborating the findings from our previous simulations^54^ using hybrid-solvent CpHMD^21^ and CHARMM PME-CpHMD.^26^ In contrast, h-bonding and solvation favor opposite microstates with the Amber force fields.

We focus on the difference between the ff14sb and ff19sb simulations to understand the switch in the preferred microstate. At pH=p*K*_1_, where the first proton binds, E35H/D52^−^ shows *N*_HB_ = 1.4 with ff14sb but 1.2 with ff19sb, indicating that D52^−^ is more stabilized by h-bonding in the ff14sb simulation (Figure 5b). At the same time, hydration of E35H/D52^−^ is slightly greater with ff14sb than ff19sb (Figure 5c). This explains why E35H/D52^−^ is the preferred state by ff14sb, whereas the alternative microstate is preferred by ff19sb. Taken together, these analyses identify local electrostatic origins of the force-field-dependent protonation of the catalytic dyad: subtle differences in the h-bond strength (ff14sb slightly stronger than ff19sb) and hydration (c22 has less hydration and slight preference for E35H/D52^−^) determine which residue binds the first proton.

### Effect of salt screening

The PME-CpHMD simulations discussed above were performed at an ionic strength of 150 mM, which is typical for MD simulations. To examine the effect of salt concentration on the calculated p*K*_a_’s, we repeated the c22 simulations of HEWL at a reduced 50 mM ionic strength (Table 3), which is the estimated experimental condition.^52^ Debye–Hückel salt screening weakens charge-charge interactions and is therefore expected to decrease the magnitude of p*K*_a_ downshifts of carboxylate residues involved in salt-bridge interactions, as demonstrated by the GB-CpHMD simulations.^55^ This is indeed the case for D66, whose p*K*_a_ is 2.3 at 150 mM and decreases to 1.7 at 50 mM ionic strength, closer to the experimental value of 1.2. However, this effect is much smaller for D48, whose p*K*_a_ remains elevated at 2.1, compared to the experimental value of 1.4.

Somewhat surprisingly, the largest effect of salt screening is on the p*K*_a_ of E35, which has an independent p*K*_a_ of 7.0 at 50 mM salt compared to 6.3 at 150 mM, enlarging the deviation from the experimental value of 6.1. In contrast, the p*K*_a_ of D52 is nearly unaffected (5.5 and 5.6 at 50 and 150 mM ionic strength, respectively). Consistent with the widened gap between the E35 and D52 p*K*_a_’s, their electrostatic coupling is decreased, making E35H/D52^−^ the predominant singly protonated state with a population of 92% (Suppl. Figure S16). Further stabilization of E35H/D52^−^ relative to E35^−^/D52H can be attributed to the increased h-bonding of D52^−^; however, the decreased solvation of E35^−^ pushes its p*K*_a_ higher (Suppl. Figure S16). Sensitivity to salt concentration is also manifested by the p*K*_a_ of H15, which decreases from 4.7 to 4.3 as the ionic strength is lowered from 150 to 50 mM, enlarging the deviation from the experimental value of 5.5. At 50 mM salt, the calculated p*K*_a_’s of most carboxylic residues (E7, D18, D66, D87, D101, D119) are brought closer to experiment, whereas the p*K*_a_ errors for the buried residues (H15 and E35) are increased. As a result, the RMSD and Pearson *r* between the experimental p*K*_a_’s and calculated values at the two salt concentrations are similar.

#### Comparison to our previous work and the GROMACS PME-CpHMD simulations

We compared the c22 HEWL p*K*_a_’s to our previous work^28^ and the GROMACS PME-CpHMD simulations with the c36m^56^ force field by Aho et al.^30^ (Table 4). 0-75 ns is used to be consistent with our previous work and Aho et al.^30^ Note, the latter used five replicates at each pH, so the actual simulation time is five times. Considering the revised model titration parameters, the p*K*_a_ differences between our present and previous work are in close agreement, with RMSD and mean unsigned deviation (MUD) of 0.28 and 0.27 units, respectively. Given the differences in the CpHMD implementation and protein force field (c22 vs. c36m^56,57^) the p*K*_a_’s from this work and Aho et al.^30^ can also be considered similar, with RMSD and MUD of 0.39 and 0.34 units, respectively, and the Pearson *r* is 0.99. A more extensive comparison between the PME-CpHMD implementations will be reported in a forthcoming work. Finally, the comparison between the p*K*_a_’s calculated from the first 75 ns and 100-150 ns shows that the p*K*_a_ of E35 is brought closer to experimental value but several depressed p*K*_a_’s of aspartic acid (D18, D48, D66, D87) move slightly away from experiment. Notably, the E35 p*K*_a_ of Aho et al. (7.6)^30^ is closer to our 0-75 ns value (7.1) than the 100-150 ns value (6.3). Therefore, these comparisons demonstrate that pH-REX PME-CpHMD simulations yield robust p*K*_a_ values with higher efficiency than independent-pH simulations.^21^

**Table 4:**
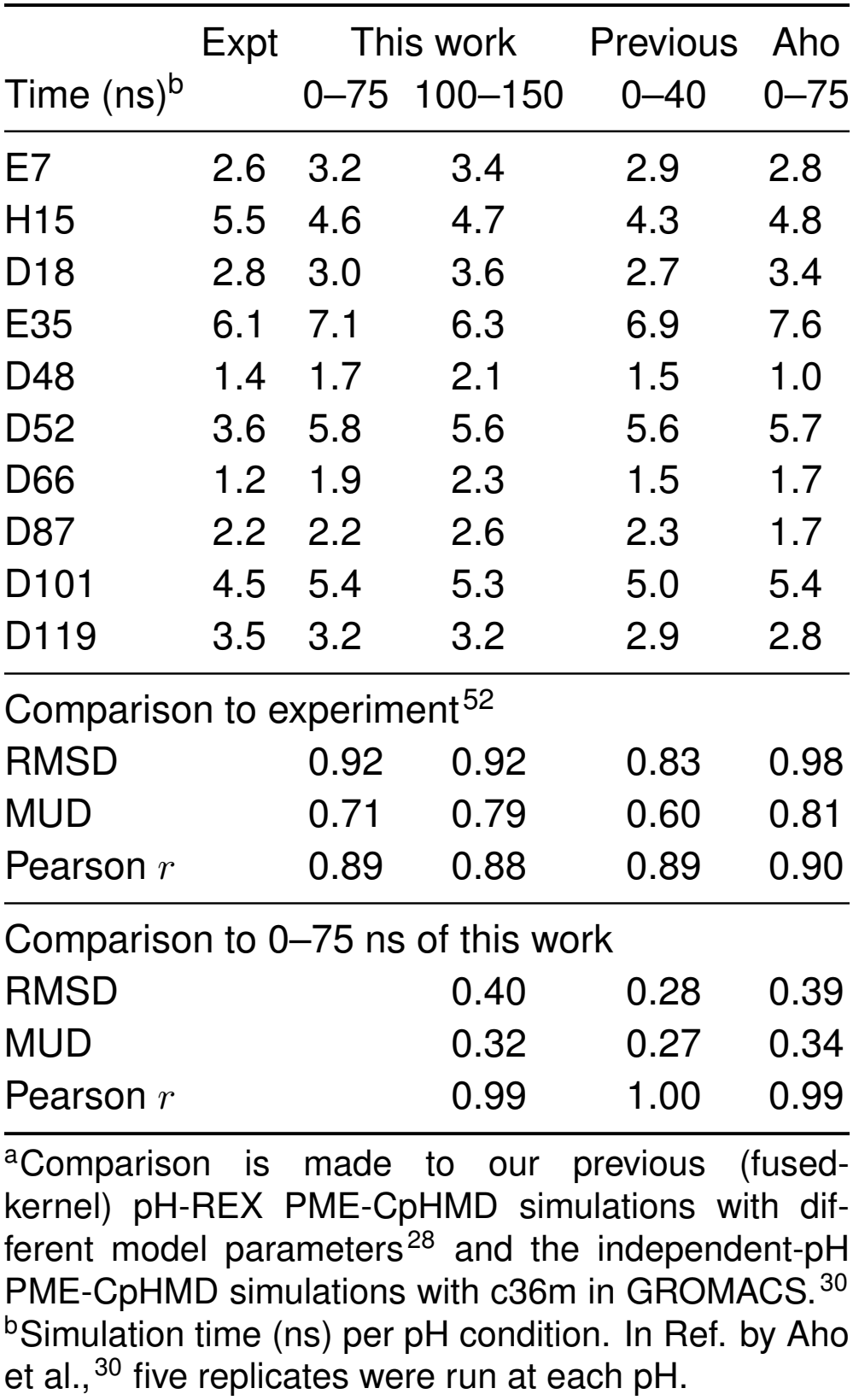
Comparison of HEWL p*K*_a_ values calculated in this work and previous studies^a^.

## 5 Conclusions

We have presented a second-generation GPU implementation of PME-CpHMD in Amber that incorporates a split-kernel nonbonded calculation, native asynchronous pH replica exchange (AREX), and revised model parameters for all six titratable residue types. Allowing titration of all Asp, Glu, His, Cys, Tyr, and Lys, the split-kernel implementation increases throughput by 1.5–2-fold over the previous implementation and keeps the overhead relative to cMD below 18% for systems of 45 to 1,338 residues or 30,000–281,000 atoms. The native AREX implementation eliminates the overhead of repeatedly stopping and restarting MD simulations in the previous wrapper-based workflow and allows replica-exchange simulations to run on fewer GPUs than pH replicas. Together, these advances substantially lower the computational barrier to AREX PME-CpHMD simulations, making them practical for routine studies of proton-coupled biomolecular processes.

Long-timescale (150 ns per replica, 2.7 µs aggregate per force field) PME-CpHMD simulations of HEWL with the c22, ff14sb, and ff19sb force fields revealed remaining challenges. Because p*K*_a_ values are a sensitive test of the microscopic description of electrostatics, deviations from experiment expose force-field limitations. Consistent with our previous simulations of the mini-protein BBL,^53^ all three force fields overestimate the p*K*_a_ downshift of H15 associated with solvent sequestration. This overestimation is modest (0.8 units) with c22 but much larger with ff14sb (3.2 units) and ff19sb (2.7 units), which may be attributed to less solvent exposure of the charged histidine in ff14sb and ff19sb simulations.

The markedly different residue-specific (independent) and stepwise p*K*_a_ values of the E35/D52 catalytic dyad across the three force fields further demonstrate how subtle differences in the electrostatic description can lead to qualitatively different biological conclusions. Specifically, D52^−^ forms weaker h-bonds with ff19sb than with ff14sb, and this small difference results in a higher p*K*_a_ value for D52 with ff19sb. As a consequence, ff19sb assigns D52 the role of proton donor, the residue that binds the first proton within a coupled titration pair. In contrast, ff14sb assigns this role to E35, with D52 serving as the nucleophile, consistent with experiment. c22 also assigns the role of proton donor to E35, but it makes both E35 and D52, particularly E35, less solvent-exposed than the Amber force fields. These results highlight the need for continued force-field development, particularly a more accurate balance between the intramolecular electrostatic interactions and solvation of charged residues. A related direction for future research is to implement PME-CpHMD with polarizable force fields, such as the Drude^58^ and AMOEBA^59^ models. A recent work has made the first step in that direction.^60^

## Supporting Information

Supporting Information is available free of charge at https://pubs.acs.org/doi/10.1021/acs.jctc.XXXXXXX.

Supplementary Methods: numerical validation of the split-kernel PME-CpHMD implementation; new and revised input controls for PME-CpHMD; and detailed parameterization procedures for six residue types.

Supplementary Tables: native AREX input controls (Table S1); new or revised PME-CpHMD phmdin controls (Table S2); updated model-parameterization protocols (Table S3); final model-potential parameters for the c22, ff14sb, and ff19sb force fields (Table S4); replica-exchange diagnostics for AREX simulations (Table S5); multi-GPU scaling performance of native AREX (Table S6); and solvent number densities and finite-size corrections (Table S7).

Supplementary Figures: thermodynamic-integration fits used to derive the model titration potentials (Figures S1–S5); time evolution of the calculated HEWL p*K*_a_ values (Figures S6–S8); titration curves of HEWL residues (Figures S9–S11); RMSD distributions of HEWL at different pH (Figure S12); distance between E35 and D52 (Figure S13); residue-specific hydrogen-bond occupancies of E35 and D52 (Figure S14); and titration as well as electrostatic environment of E35/D52 from the c22 simulations at 50 mM ionic strength (Figure S16) (PDF).

## Acknowledgments

Financial support by the National Institutes of Health (R35GM148261) is acknowledged.

G.W.D. thanks Dr. Mingzhe Shen for providing the ASIC system used in the performance benchmarks, Dr. Subarna Sasmal for providing the TacF system used in the performance benchmarks, and Dr. Aarion Romany for technical guidance on molecular-system preparation. ChatGPT (OpenAI) and Claude (Anthropic) were used for language editing and manuscript organization. The authors reviewed and edited all content and take full responsibility for the manuscript.

## Associated Content

### Data and Software Availability Statement

All simulation inputs, starting structures, parameterization and benchmark data, the numerical data underlying the figures, and the analysis and fitting scripts are available on Zenodo at https://doi.org/10.5281/zenodo.22003475. The CpHMD model parameter files for ff19SB, ff14SB, and c22 are available on Zenodo at https://doi.org/10.5281/zenodo.22003318.

The split-kernel PME-CpHMD and native AREX implementations were developed in the Amber24 pmemd.cuda and pmemd.cuda.MPI codebase. The source-code patch is available on Zenodo (https://doi.org/10.5281/zenodo.21997770). The patch and the CpHMD model parameter files for the c22, ff14sb, and ff19sb force fields will be deposited in Amber.

## Notes

### Competing Interest Statement

The authors have declared no competing interest.

