## Supporting Information for "High-Performance All-Atom Continuous Constant-pH Molecular Dynamics and Native Asynchronous Replica Exchange in Amber"

### Contents

**List of Tables** 2

**List of Figures** 3

**Supplementary Methods** 4

**1 Numerical validation of the split-kernel PME-CpHMD implementation** 4

**2 Updated input controls for PME-CpHMD and AREX** 5

**3 Parameterization of model titration potentials** 5

**Supplementary Tables** 12

**Supplementary Figures** 18

**References** 33

S7 Solvent number densities and finite-size corrections for HEWL simulations . 16
S8 Comparison of HEWL $\mathrm{p}K_\mathrm{a}$ values calculated in this work and previous studies 17

### Supplementary Methods

#### 1 Numerical validation of the split-kernel PME-CpHMD implementation

The split-kernel direct-space PME-CpHMD implementation, in which the correction terms are accumulated into the existing force, virial, energy, and titration-coordinate derivative accumulators, was validated against the previous fused-kernel implementation retained in guarded validation builds. Both were evaluated with identical coordinates, titration variables, neighbor lists, and nonbonded parameters, and all four accumulators were compared for the force-only, force-switching, virial, and energy/virial execution paths.

The two implementations agreed within the tolerances expected for the accumulator format. Maximum component-wise force differences were typically $10^{-3}$–$10^{-2}$ kcal mol$^{-1}$ Å$^{-1}$; with the Amber fixed-point force scale of $2^{32}$, the strict tolerance of $2^{24}$ accumulator units corresponds to approximately $4 \times 10^{-3}$ kcal mol$^{-1}$ Å$^{-1}$. In the force-switching path, the observed maximum differences (0.006–0.008 kcal mol$^{-1}$ Å$^{-1}$) were well below the looser tolerance of 0.0625 kcal mol$^{-1}$ Å$^{-1}$. Reduced titration-coordinate derivatives differed by approximately $10^{-5}$, van der Waals energies by approximately $10^{-7}$ kcal mol$^{-1}$, and direct-space electrostatic energies not at all. The largest virial-component differences, approximately 0.25 kcal mol$^{-1}$ in strict force-switching edge cases, correspond to pressure differences well below the instantaneous pressure fluctuations in constant-pressure MD.

These residual differences arise from the order of fixed-point conversion and accumulation: the fused path effectively accumulates $\mathrm{round}(x_{\mathrm{PME}} + x_{\mathrm{CpHMD}})$, whereas the split path accumulates $\mathrm{round}(x_{\mathrm{PME}}) + \mathrm{round}(x_{\mathrm{CpHMD}})$. The two are algebraically equivalent but need not be bitwise identical, so the discrepancies reflect numerical ordering, not a change in the CpHMD potential.

### 2 Updated input controls for PME-CpHMD and AREX

**Updated PME-CpHMD `phmdin` controls.** The `phmdin` namelist now supports fixed-coordinate thermodynamic integration (TI) and an extended residue-selection syntax for titratable sites (Table S2).

**Native AREX controls.** Native asynchronous replica exchange (AREX) is configured through the standard `&cntrl` namelist (Table S1); the currently implemented exchange type is pH replica exchange (`phrex`). Note, Tables S1 and Table S2 list only the input controls that are new or revised in this work.

### 3 Parameterization of model titration potentials

In continuous CpHMD,[1,2] the protonation and tautomer coordinates are expressed as $\lambda = \sin^2\theta$ and $x = \sin^2\theta_x$, respectively, where $\theta$ and $\theta_x$ are the dynamic variables propagated in the simulation. Parameterization of model titration potentials was performed in the angular coordinates $\theta$ and $\theta_x$ rather than in $\lambda$ and $x$, which minimizes error propagation.[3] The mean forces, $\langle \partial U/\partial\theta \rangle$ and $\langle \partial U/\partial\theta_x \rangle$, were calculated from TI simulations of the blocked tripeptides. The mean forces were fit by unweighted least squares, giving equal weight to each fixed-coordinate condition. Compared with previous work,[2,3] the updated protocol samples more $\theta$ (and $\theta_x$) values for all residue types and, for double-site titrations, fits the mean forces directly to the analytical forms of the two-dimensional titration potentials (Table S3). Because $\mathrm{d}\lambda/\mathrm{d}\theta = \sin 2\theta$ vanishes at $\theta = 0$ and $\pi/2$ (and likewise for $\theta_x$), the generalized-force component associated with a coordinate at one of these endpoints was set to its analytical value of zero.

#### 3.1 Parameterization for carboxylic acids

Asp and Glu titrations are represented using the double-site model,[2] in which $\theta$ and $\theta_x$ represent protonation and tautomer coordinates, respectively. Using TI simulations, the mean forces on a two-dimensional grid were calculated,

$$\theta \in \{0.0,\ 0.3,\ 0.4,\ 0.6,\ 0.7854,\ 1.0,\ 1.2,\ 1.3,\ 1.4\}, \tag{1}$$

$$\begin{aligned}\theta_x \in \{&0.0,\ 0.1,\ 0.3,\ 0.4,\ 0.5,\ 0.6,\ 0.7854,\ 1.0,\ 1.1,\\ &1.2,\ 1.3,\ 1.4,\ 1.5,\ 1.5708\},\end{aligned} \tag{2}$$

where 0.7854 and 1.5708 correspond to $\pi/4$ and $\pi/2$, respectively. The final fitting grid contained 126 ($9 \times 14$) coordinate pairs. TI simulations were performed at 124 of these points. The two corner points, $(\theta, \theta_x) = (0, 0)$ and $(0, \pi/2)$, for which both generalized-force components vanish analytically, were supplied as zero-force boundary values without simulation. At other endpoint-containing grid points, only the generalized-force component associated with the endpoint coordinate vanishes; the orthogonal component was obtained from the TI simulation.

The parameters were obtained by staged fitting. At each fixed $\theta$, the mean forces along $\theta_x$ coordinate were fit to the derivative of a quadratic function,[2]

$$\left\langle \frac{\partial U}{\partial \theta_x} \right\rangle_\theta = 2A(\theta)\sin(2\theta_x)\left[\sin^2\theta_x - B(\theta)\right], \tag{3}$$

yielding $A(\theta)$ and $B(\theta)$ (upper rows of Figures S1 and S2). Because $\theta = \pi/2$ was not included in the simulated carboxylate grid, a zero-force point at this endpoint was added analytically to each fixed-$\theta_x$ fit as a boundary condition.

Similarly, at each fixed $\theta_x$, the mean forces along the $\theta$ coordinate were fit to

$$\left\langle \frac{\partial U}{\partial \theta} \right\rangle_{\theta_x} = 2A(\theta_x)\sin(2\theta)\left[\sin^2\theta - B(\theta_x)\right], \tag{4}$$

yielding $A(\theta_x)$ and $B(\theta_x)$ (lower rows of Figures S1 and S2).

The values of $A(\theta)$ obtained from Equation 3 were then fit to a second-order polynomial,[2]

$$A(\theta) = R_1 \sin^4 \theta + R_2 \sin^2 \theta + R_3, \tag{5}$$

yielding $R_1$, $R_2$, and $R_3$ (Figure S3). Although $R_4$ can be estimated as the mean of the fitted $B(\theta)$ values, it was fixed at 0.5, reflecting the equivalence of the two carboxylate oxygens.

The carboxylate model is conventionally expressed in terms of six parameters, $R_1$–$R_6$,[2] and the AMBER CpHMD parameter file likewise contains six corresponding entries.[4] Before evaluating the potential, however, the AMBER implementation algebraically transforms the fifth and sixth entries. To distinguish the resulting internal coefficients from the conventional $R_1$–$R_6$ notation, we denote them here as $C_5$ and $C_6$:

$$\begin{aligned} C_5 &= R_5 - R_1 R_4^2, \\ C_6 &= -2R_5R_6 - R_2R_4^2. \end{aligned} \tag{6}$$

The model potential evaluated by AMBER is therefore

$$U_{\text{mod}}(\lambda, x) = \left(R_1\lambda^2 + R_2\lambda + R_3\right)(x - R_4)^2 + C_5\lambda^2 + C_6\lambda. \tag{7}$$

At fixed $\theta_x$, with $x = \sin^2 \theta_x$, comparison of Equation 7 with the fitted quadratic $A(\theta_x)[\lambda - B(\theta_x)]^2$ gives the slice-wise estimates

$$C_5(\theta_x) = A(\theta_x) - R_1 \left[\sin^2 \theta_x - R_4\right]^2 \tag{8}$$

and

$$C_6(\theta_x) = -2A(\theta_x)B(\theta_x) - R_2 \left[\sin^2 \theta_x - R_4\right]^2. \tag{9}$$

The final $C_5$ and $C_6$ coefficients were taken as the arithmetic means of the corresponding values over all fixed-$\theta_x$ slices.

This differs from the previous parameterization protocol,[3] in which $R_5$ and $R_6$ were obtained through separate secondary fits of $A(\theta_x)$ and $B(\theta_x)$, respectively. Here, once $R_1$–$R_4$ have been determined, $C_5$ and $C_6$ are obtained directly from the analytical relationships imposed by the implemented two-dimensional potential. This eliminates those two secondary fits and provides independent estimates of the same coefficients from each fixed-$\theta_x$ slice.

Finally, the fifth and sixth CpHMD parameter-file entries were recovered by inverting Equation 6,

$$R_5 = C_5 + R_1 R_4^2, \tag{10}$$

$$R_6 = -\frac{C_6 + R_2 R_4^2}{2R_5}. \tag{11}$$

The resulting parameter-file values $R_1$–$R_6$ are listed in Table S4.

#### 3.2 Parameterization for histidine

His titration is represented by a double-site model with two nonequivalent microscopic pathways.[2] The TI grid comprised three intersecting one-dimensional branches: two protonation branches, in which $\theta$ was varied at fixed $\theta_x = 0$ and $\pi/2$ over

$$\theta \in \{0.0,\ 0.1,\ 0.2,\ 0.4,\ 0.5,\ 0.6,\ 0.7,\ 0.7854,\ 0.9,\ 1.0,\ 1.1,\ 1.2,\ 1.3,\ 1.4,\ 1.5,\ 1.5708\}, \tag{12}$$

and a neutral-tautomer branch, in which $\theta_x$ was varied at fixed $\theta = \pi/2$ over

$$\theta_x \in \{0.0,\ 0.1,\ 0.2,\ 0.3,\ 0.4,\ 0.5,\ 0.6,\ 0.7,\ 0.7854,\ 1.0,\ 1.1,\ 1.2,\ 1.3,\ 1.4,\ 1.5,\ 1.5708\}. \tag{13}$$

Counting the two shared intersection points once, this gives 46 simulations.

Although the three sampled branches are one-dimensional profiles, they were not fit independently. Instead, the generalized-force data from all three branches were combined in a single global least-squares fit to the derivatives of one common two-dimensional model potential,[2]

$$U_{\text{mod}}(\lambda, x) = A_{10}\lambda^2 x^2 + 2(C_1 - C_0)\lambda x - 2C_1\lambda \\ + 2(C_0 - C_1 - C_{10})\lambda^2 x + A_1\lambda^2, \tag{14}$$

where

$$C_1 = A_1 B_1, \qquad C_0 = A_0 B_0, \qquad C_{10} = A_{10} B_{10}. \tag{15}$$

On the three sampled boundary branches, the model reduces to one-dimensional generalized-force expressions. At $x = 0$,

$$\left\langle \frac{\partial U}{\partial \theta} \right\rangle_{x=0} = 2\sin(2\theta)\left[A_1 \sin^2\theta - C_1\right], \tag{16}$$

whereas at $x = 1$,

$$\left\langle \frac{\partial U}{\partial \theta} \right\rangle_{x=1} = 2\sin(2\theta)\left[A_0 \sin^2\theta - C_0\right], \tag{17}$$

with

$$A_0 = A_1 + A_{10} + 2(C_0 - C_1 - C_{10}). \tag{18}$$

Along the neutral-tautomer branch, $\lambda = 1$,

$$\left\langle \frac{\partial U}{\partial \theta_x} \right\rangle_{\lambda=1} = 2\sin(2\theta_x)\left[A_{10} \sin^2\theta_x - C_{10}\right]. \tag{19}$$

All generalized-force values from these three profiles were included in a single fit with the shared parameter vector

$$[A_1,\ C_1,\ C_0,\ A_{10},\ C_{10}]. \tag{20}$$

Thus, "one-dimensional" refers to the sampled boundary profiles, not to separate parameter fits. Because the orthogonal generalized force vanishes identically on each boundary branch, only $\partial U/\partial\theta$ contributes on the two protonation branches and only $\partial U/\partial\theta_x$ contributes on the neutral-tautomer branch.

The fitted quantities were converted to the six CpHMD parameter-file values according to

$$B_1 = \frac{C_1}{A_1}, \tag{21}$$

$$B_{10} = \frac{C_{10}}{A_{10}}, \tag{22}$$

$$A_0 = A_1 + A_{10} + 2(C_0 - C_1 - C_{10}), \tag{23}$$

$$B_0 = \frac{C_0}{A_0}. \tag{24}$$

Accordingly, $(A_1, B_1)$ describes the $x = 0$ protonation branch, $(A_0, B_0)$ the $x = 1$ protonation branch, and $(A_{10}, B_{10})$ the $\lambda = 1$ neutral-tautomer branch (Table S4 and Figure S4).

#### 3.3 Parameterization for single-site model titration

Cys, Tyr, and Lys were represented using the single-site model titration, with TI simulations at

$$\theta \in \{0.0,\ 0.1,\ 0.2,\ 0.3,\ 0.4,\ 0.5,\ 0.6,\ 0.7854,\ 1.0,\ 1.1,\ 1.2,\ 1.3,\ 1.4,\ 1.5,\ 1.5708\}, \tag{25}$$

giving 15 simulations for each residue and force field. The model potential

$$U_{\text{mod}}(\lambda) = A(\lambda - B)^2, \tag{26}$$

has the angular generalized force

$$\frac{\partial U_{\mathrm{mod}}}{\partial \theta} = 2A \sin(2\theta) \left[\sin^2 \theta - B\right], \tag{27}$$

to which the mean generalized forces were fit directly to obtain $A$ and $B$ (Table S4 and Figure S5).

### Supplementary Tables

Table S1: Native AREX input controls in the `&cntrl` namelist.

| Control | Default | Function |
|---|---|---|
| `arex` | 0 | Enables native AREX when set to `1`. |
| `arex_type` | blank | Selects the exchange type; presently `'phrex'` for pH-AREX. |
| `arex_segment_steps` | 0 | Number of MD steps between the exchange attempts (one AREX segment or cycle). `nstlim` must be an exact multiple of this value. |
| `arex_warmup_segments` | 0 | Number of AREX segments propagated for each replica before exchanges begin. |
| `arex_num_workers` | 0 | Number of AREX workers (GPUs). A value of `0` uses all available MPI ranks except the dedicated global-scheduler rank; a positive value explicitly sets the number of workers. |
| `arex_ladder` | unset | Ordered exchange-coordinate ladder. For `phrex`, this is the pH ladder, for example: 5, 6, 7, 8. At least two entries are required. |
| `arex_seed` | 0 | Controls initialization of the AREX random-number stream. A value of `0` retains the default random-number state, a positive value specifies a reproducible seed, and a negative value requests clock-based seeding. |
| `arex_chkfrq` | 0 | Number of completed AREX segments between periodic checkpoint writes. A value of `0` disables periodic checkpointing. A final checkpoint is always written when the AREX campaign completes, allowing the calculation to be continued. |
| `arex_emulate_restart` | 0 | Selects segment-boundary handling. The default (`0`) runs AREX segments continuously; `1` reproduces the stop/start segment-boundary behavior of the previous Python-based workflow. |
| `arex_restraint_npoints`, `arex_restraint_segment`, `arex_restraint_force` | disabled | Optional positional-restraint schedule during AREX warmup. Setting `arex_restraint_npoints=0` disables the schedule. Each `arex_restraint_segment` value is a zero-based completed-slot-segment boundary, and the corresponding `arex_restraint_force` value applies from that boundary until the next one; the final value persists. The schedule requires positional restraints (`ntr=1`) and a positive number of AREX warmup segments. |

Table S2: New or revised PME-CpHMD `phmdin` controls.[a]

| Control | Default | Function |
|---|---|---|
| `qphmd_ti` | `.false.` | Enables the fixed-coordinate thermodynamic-integration mode. When enabled, `phtest_theta` is required and the implementation sets `phtest=1`, `prderiv=.true.`, `prlam=.false.`, `prdudls=.false.`, and `qphmdstart=.true.`; the ordinary `phmdrestrt` input is not read. |
| `phtest_theta` | unset | Specifies the fixed angular titration-coordinate value or values used in `qphmd_ti` mode. The first value is required, and multiple coordinates are supplied as a contiguous sequence for systems containing more than one titration coordinate. |
| `masktitrres` | blank | Selects residues for PME-CpHMD titration. A residue name alone, for example `CYS`, selects all residues of that type; `CYS:42` selects residue number 42 specifically; and `CYS:!42` selects all Cys residues except residue 42. Repeated indexed selectors can define inclusion or exclusion lists for a residue type. |

[a] `masktitrrestypes` is deprecated: it is still parsed for backward compatibility but is ignored. Active titratable residue types are inferred from the nonblank `masktitrres` selectors.

Table S3: Comparison of the previous[2,3] and updated protocols for model-potential parameterization.

| Aspect | Previous[2,3] | Updated | Rationale or consequence |
|---|---|---|---|
| TI setup | $\theta$ or $\theta_x$ values supplied through the CpHMD restart-state workflow | $\theta$ or $\theta_x$ values specified through `phmdin` | Eliminates manual editing of the restart files and simplifies TI setup. |
| Number of TI runs | Limited $\theta$ or $\theta_x$ conditions | Denser $\theta$ and $\theta_x$ grids for all residue types | Broader sampling of the mean-force surfaces for more accurate fitting. |
| Asp/Glu fitting | Separate secondary fits of $A(\theta_x)$ and $B(\theta_x)$ to obtain the final two carboxylate parameters | Internal coefficients $C_5$ and $C_6$ obtained directly from the analytical relationships of the implemented two-dimensional potential for each fixed-$\theta_x$ slice and averaged across slices | Eliminates two secondary fits and provides independent estimates of the same coefficients across the fixed-$\theta_x$ slices. |
| His fitting | Fitting to one-dimensional mean-force profiles | All three generalized-force profiles fit simultaneously to a single two-dimensional model | Constrains all sampled branches to a common underlying model potential. |
| Titr. residue types | Asp, Glu, Cys, His, and Lys | Asp, Glu, Cys, His, Lys, and Tyr | Tyr added. |

Table S4: Final model-potential parameters used for PME-CpHMD simulations with c22, ff14sb, and ff19sb force fields.[a]

| Residue | $R_1$ | $R_2$ | $R_3$ | $R_4$ | $R_5$ | $R_6$ |
|---|---|---|---|---|---|---|
| **c22** | | | | | | |
| Asp | –33.395204 | 77.219141 | –43.733904 | 0.500000 | –74.611641 | 0.104421 |
| Glu | –33.726186 | 77.516003 | –43.747875 | 0.500000 | –74.456661 | 0.075401 |
| His | –52.161892 | 0.373673 | –52.147355 | 0.518370 | –45.425080 | 0.333849 |
| Cys | –82.039571 | –0.004798 | N/A | N/A | N/A | N/A |
| Tyr | –70.512644 | –0.437396 | N/A | N/A | N/A | N/A |
| Lys | –72.548258 | 0.641759 | N/A | N/A | N/A | N/A |
| **ff14sb** | | | | | | |
| Asp | –40.774211 | 89.873660 | –49.236443 | 0.500000 | –46.827874 | 0.217352 |
| Glu | –39.922736 | 89.011950 | –49.158969 | 0.500000 | –45.282403 | 0.180961 |
| His | –34.790976 | 0.246319 | –33.321952 | 0.206225 | –33.143444 | 0.529066 |
| Cys | –77.999644 | –0.042434 | N/A | N/A | N/A | N/A |
| Tyr | –30.191213 | –0.327209 | N/A | N/A | N/A | N/A |
| Lys | –52.943024 | 0.489999 | N/A | N/A | N/A | N/A |
| **ff19sb** | | | | | | |
| Asp | –38.405279 | 87.560932 | –49.316716 | 0.500000 | –49.483708 | 0.184914 |
| Glu | –37.679381 | 86.941144 | –49.378055 | 0.500000 | –48.160014 | 0.149072 |
| His | –35.015469 | 0.212275 | –33.026611 | 0.167217 | –33.324377 | 0.527483 |
| Cys | –82.697182 | –0.039967 | N/A | N/A | N/A | N/A |
| Tyr | –32.063688 | –0.341620 | N/A | N/A | N/A | N/A |
| Lys | –54.122864 | 0.465016 | N/A | N/A | N/A | N/A |

[a]For Asp and Glu, the six columns correspond to $R_1$–$R_6$. For His, they correspond to $A_0$, $B_0$, $A_1$, $B_1$, $A_{10}$, and $B_{10}$, respectively. For Cys, Tyr, and Lys, the first two columns correspond to $A$ and $B$. All values are the quantities written to the Amber CpHMD parameter files. For Asp and Glu, the fifth and sixth parameter-file entries are $R_5$ and $R_6$; AMBER converts them to the internal coefficients $C_5$ and $C_6$ defined above before evaluating the model potential. For Asp and Glu, $R_4 = 0.5$ was imposed by symmetry. Energy-valued coefficients are in kcal mol$^{-1}$; $R_4$, $R_6$, and all $B$ coefficients are dimensionless.

Table S5: Replica-exchange diagnostics for pH-REX simulations of the Asp model peptide run on 1–4 GPUs. [a]

| Number of GPUs | Calculated $\mathrm{p}K_\mathrm{a}$ | Overall exchange acceptance (%) | Round trips per replica | Full-ladder visitation |
|---|---|---|---|---|
| 1 | $3.897 \pm 0.051$ | $80.90 \pm 0.20$ | $252.5 \pm 15.7$ | $100\%$ |
| 2 | $3.909 \pm 0.057$ | $80.98 \pm 0.16$ | $259.7 \pm 12.3$ | $100\%$ |
| 3 | $3.897 \pm 0.041$ | $81.10 \pm 0.19$ | $263.4 \pm 12.5$ | $100\%$ |
| 4 | $3.902 \pm 0.053$ | $80.98 \pm 0.14$ | $258.1 \pm 9.9$ | $100\%$ |

[a] Ten independent simulations of the ff19sb Asp model peptide were performed with each of one to four AREX workers (GPUs). Each simulation used six pH conditions (2.5–5.0 in increments of 0.5), with 21 ns of sampling per condition and exchange attempts every 1000 MD steps (2 ps). The first 1 ns was excluded from analysis. All other simulation settings and the $\mathrm{p}K_\mathrm{a}$ fitting procedure were identical across worker counts. Exchange acceptance is the fraction of attempted exchanges that were accepted. Round trips (pH 2.5 $\rightarrow$ 5.0 $\rightarrow$ 2.5) were averaged over the six replicas of each simulation. Values are means $\pm$ standard deviations over ten simulations. Full-ladder visitation is the percentage of the 60 replica trajectories per worker count that visited all six pH conditions.

Table S6: Scaling performance of native AREX across four protein systems.[a]

| | | AREX progress rate (ns day$^{-1}$) | | | | Four-worker scaling | | Four-worker aggregate rate |
|---|---|---|---|---|---|---|---|---|
| System | Atoms | 1 worker | 2 workers | 3 workers | 4 workers | Speedup | Efficiency (%) | ns day$^{-1}$ |
| BBL | 29,875 | 9.86 | 19.29 | 28.27 | 36.28 | 3.68 | 92.0 | 870.7 |
| HEWL | 45,083 | 8.00 | 15.76 | 22.78 | 28.89 | 3.61 | 90.3 | 693.4 |
| TacF[b] | 130,480 | 4.45 | 8.63 | 12.53 | 15.84 | 3.56 | 89.0 | 380.2 |
| ASIC trimer[b] | 281,354 | 2.57 | 5.13 | 7.56 | 9.57 | 3.72 | 93.1 | 229.7 |

[a]Each benchmark used 24 pH slots and advanced every slot by 2 ns. AREX progress rate is the common per-pH simulation time divided by wall-clock time; speedup and parallel efficiency are relative to the one-worker rate, and the aggregate rate is 24 times the four-worker progress rate. [b]Transmembrane system.

Table S7: Solvent number densities and finite-size corrections for the HEWL $\mathrm{p}K_\mathrm{a}$ calculations.[a]

| Force field | Reference | Model-peptide density (Å$^{-3}$) | HEWL density (Å$^{-3}$) | Density difference (Å$^{-3}$) | Conversion factor (Å$^3$) | $\Delta \mathrm{p}K_\mathrm{a,corr}$ |
|---|---|---|---|---|---|---|
| | Asp | 0.033 480 | 0.032 427 | −0.001 053 | 387.0739 | −0.40 |
| c22 | Glu | 0.033 471 | 0.032 427 | −0.001 044 | 387.0739 | −0.40 |
| | His | 0.033 466 | 0.032 427 | −0.001 039 | 387.0739 | −0.40 |
| | Asp | 0.032 771 | 0.031 794 | −0.000 977 | 387.0739 | −0.37 |
| ff14sb | Glu | 0.032 763 | 0.031 794 | −0.000 969 | 387.0739 | −0.37 |
| | His | 0.032 753 | 0.031 794 | −0.000 959 | 387.0739 | −0.37 |
| | Asp | 0.033 192 | 0.032 124 | −0.001 068 | 506.2317 | −0.54 |
| ff19sb | Glu | 0.033 184 | 0.032 124 | −0.001 061 | 506.2317 | −0.54 |
| | His | 0.033 173 | 0.032 124 | −0.001 050 | 506.2317 | −0.54 |

[a] Solvent number densities were calculated as $\rho = \langle N_\mathrm{water}/V \rangle$, where $N_\mathrm{water}$ is the number of water molecules and $V$ is the instantaneous simulation box volume. For HEWL, densities were averaged over the production trajectories at all 18 pH conditions. For each model peptide, densities were averaged over the retained 20 ns at all pH conditions within each independent simulation, then averaged across three simulations. The correction was calculated as[4,5] $\Delta \mathrm{p}K_\mathrm{a,corr} = C(\rho_\mathrm{HEWL} - \rho_\mathrm{model})$, where $C = 387.0739$ Å$^3$ for CHARMM-modified and standard TIP3P and $C = 506.2317$ Å$^3$ for OPC, obtained with $T = 300$ K, $\gamma^\mathrm{s} = 0.764\, e\,$Å$^2$ for the TIP3P models, and $\gamma^\mathrm{s} = 0.9991916\, e\,$Å$^2$ for OPC.

Table S8: Comparison of HEWL p$K_a$ values calculated in this work and previous studies[a]

| | Expt | This work | | Previous | Aho |
|---|---|---|---|---|---|
| Time (ns)[b] | | 0–75 | 100–150 | 0–40 | 0–75 |
| E7 | 2.6 | 3.2 | 3.4 | 2.9 | 2.8 |
| H15 | 5.5 | 4.6 | 4.7 | 4.3 | 4.8 |
| D18 | 2.8 | 3.0 | 3.6 | 2.7 | 3.4 |
| E35 | 6.1 | 7.1 | 6.3 | 6.9 | 7.6 |
| D48 | 1.4 | 1.7 | 2.1 | 1.5 | 1.0 |
| D52 | 3.6 | 5.8 | 5.6 | 5.6 | 5.7 |
| D66 | 1.2 | 1.9 | 2.3 | 1.5 | 1.7 |
| D87 | 2.2 | 2.2 | 2.6 | 2.3 | 1.7 |
| D101 | 4.5 | 5.4 | 5.3 | 5.0 | 5.4 |
| D119 | 3.5 | 3.2 | 3.2 | 2.9 | 2.8 |
| Comparison to experiment[6] | | | | | |
| RMSD | | 0.92 | 0.92 | 0.83 | 0.98 |
| MUD | | 0.71 | 0.79 | 0.60 | 0.81 |
| Pearson $r$ | | 0.89 | 0.88 | 0.89 | 0.90 |
| Comparison to 0–75 ns of this work | | | | | |
| RMSD | | | 0.40 | 0.28 | 0.39 |
| MUD | | | 0.32 | 0.27 | 0.34 |
| Pearson $r$ | | | 0.99 | 1.00 | 0.99 |

[a]Comparison is made to our previous (fused-kernel) pH-REX PME-CpHMD simulations with different model parameters[4] and the independent-pH PME-CpHMD simulations with c36m in GROMACS.[7]
[b]Simulation time (ns) per pH condition. In Ref. by Aho et al.,[7] five replicates were run at each pH.

### Supplementary Figures

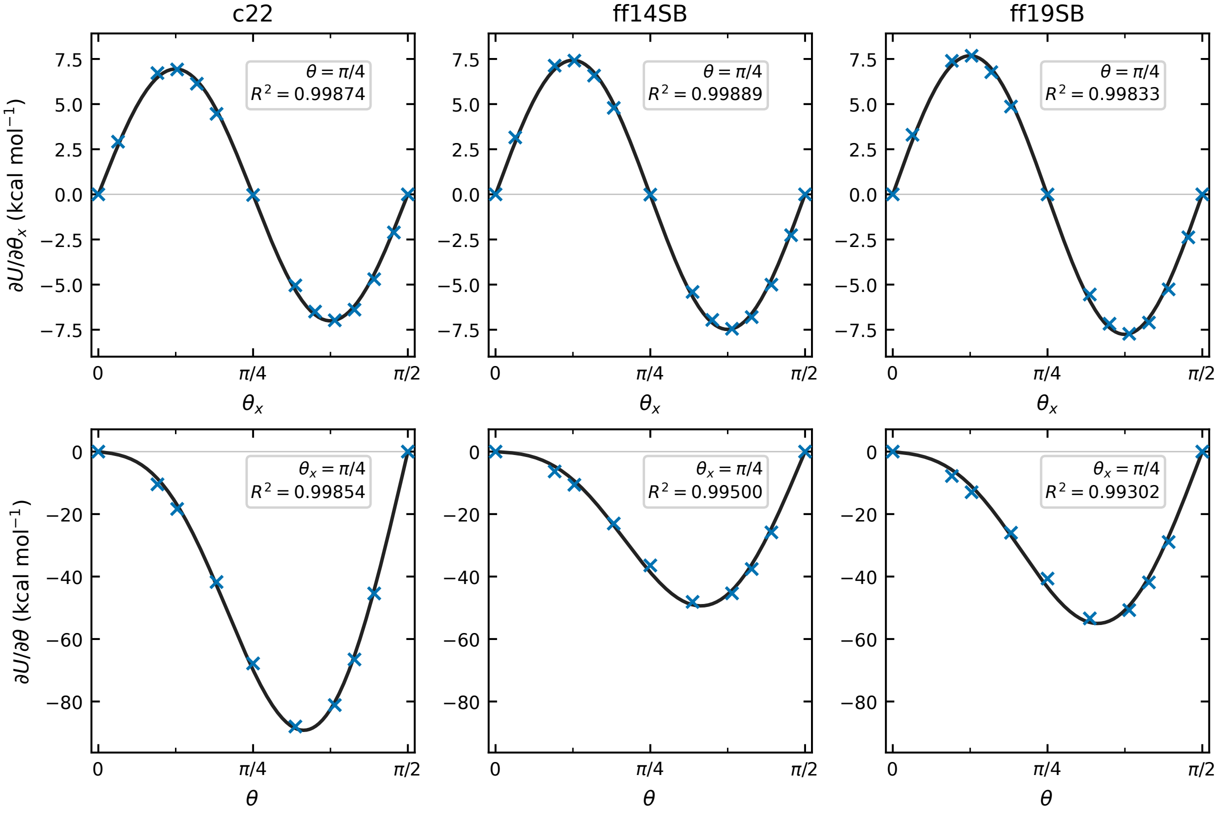

Figure S1: **Representative first-stage thermodynamic-integration fits for the Asp model peptide.** The upper row shows $\partial U/\partial\theta_x$ as a function of $\theta_x$ at fixed $\theta = \pi/4$, and the lower row shows $\partial U/\partial\theta$ as a function of $\theta$ at fixed $\theta_x = \pi/4$. Columns correspond, from left to right, to c22, ff14sb, and ff19sb. Symbols represent the generalized-force values used in the fits, and solid curves represent the fitted functions; the $\theta = \pi/2$ point in each lower-row panel is the zero-force boundary condition. Common vertical-axis limits are used within each row.

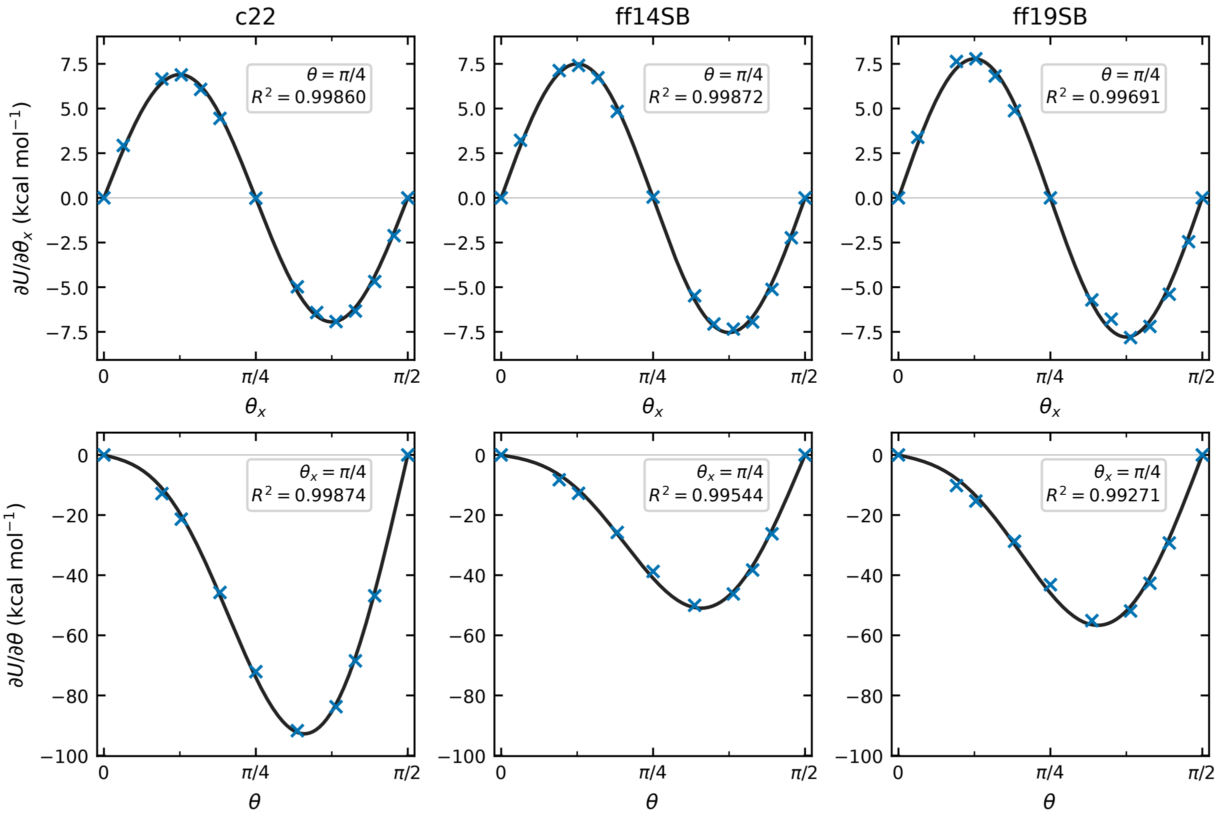

Figure S2: **Representative first-stage thermodynamic-integration fits for the Glu model peptide.** Panels and symbols are as in Figure S1.

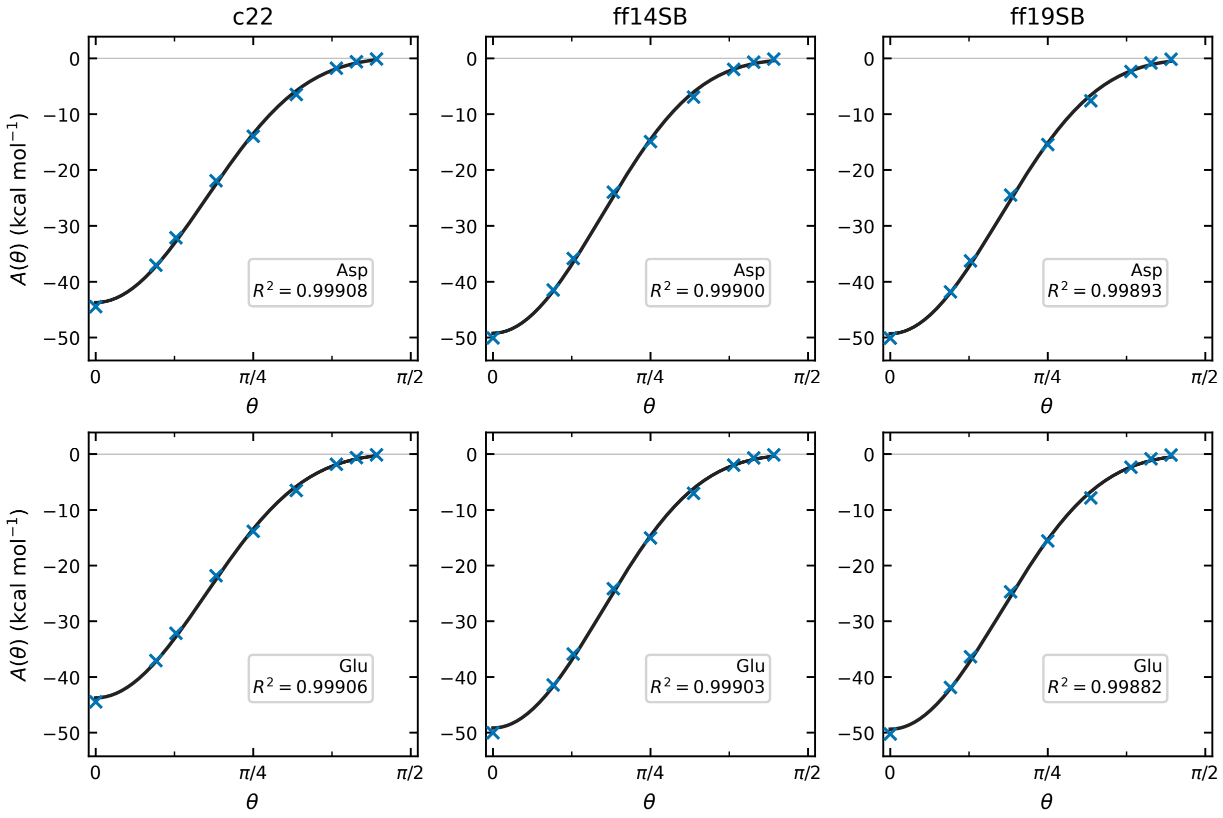

Figure S3: **Secondary fits used to determine the $R_1$, $R_2$, and $R_3$ parameters for the Asp and Glu model peptides.** Rows correspond, from top to bottom, to Asp and Glu, and columns correspond, from left to right, to c22, ff14sb, and ff19sb. Symbols represent the $A(\theta)$ values obtained from the first-stage fixed-$\theta$ fits, and solid curves represent fits to $A(\theta) = R_1 \sin^4\theta + R_2 \sin^2\theta + R_3$. Common vertical-axis limits are used across all panels.

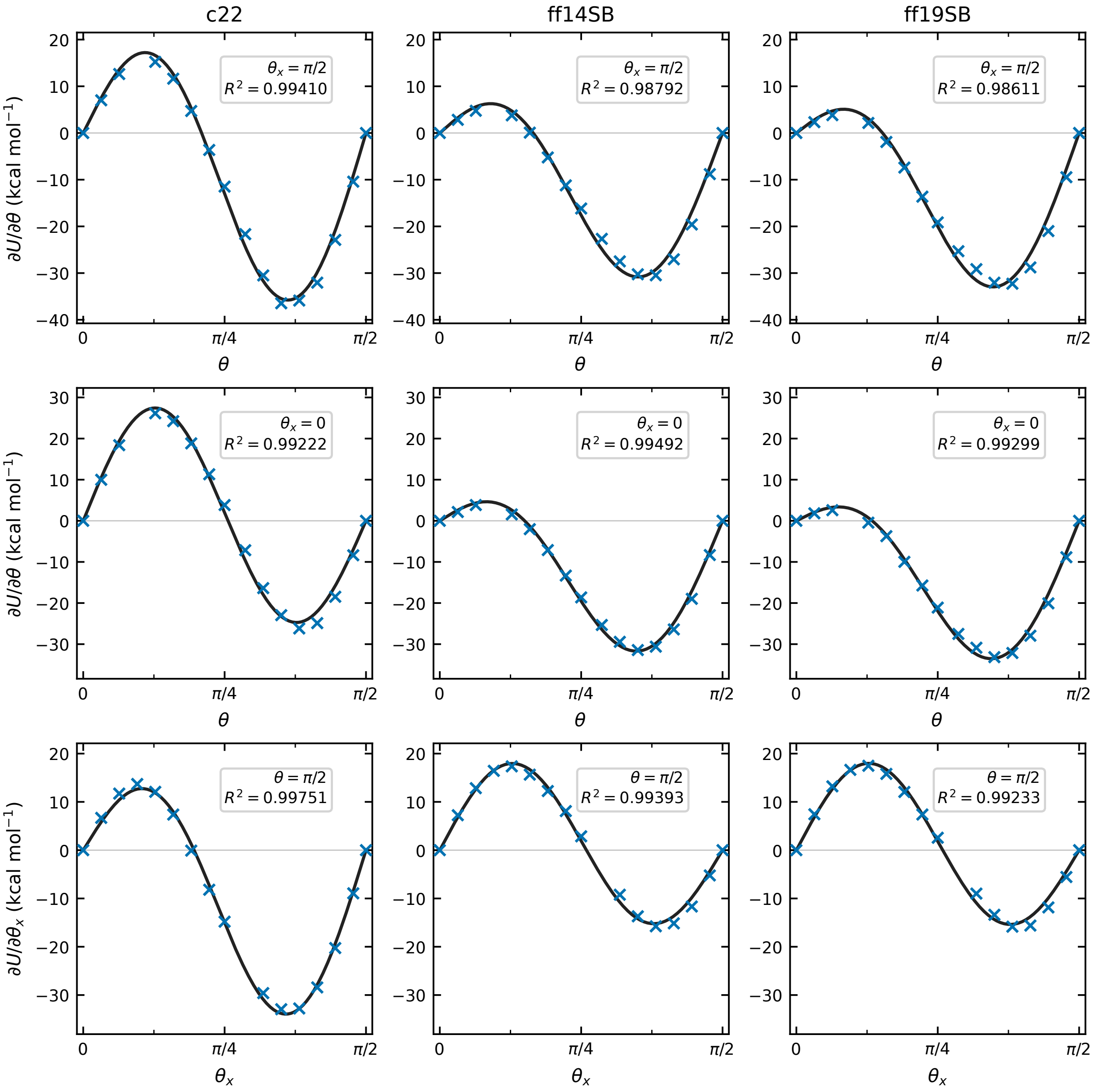

Figure S4: **Representative $\theta$-force profiles from the global model-potential fits for the His model peptide.** The upper row shows $\partial U/\partial\theta$ as a function of $\theta$ at fixed $\theta_x = \pi/2$, the middle row shows $\partial U/\partial\theta$ as a function of $\theta$ at fixed $\theta_x = 0$, and the lower row shows $\partial U/\partial\theta_x$ as a function of $\theta_x$ at fixed $\theta = \pi/2$. Columns correspond, from left to right, to c22, ff14sb, and ff19sb. Symbols represent the generalized-force values used in the global fit, and solid curves represent predictions from the final global model. Common vertical-axis limits are used within each row.

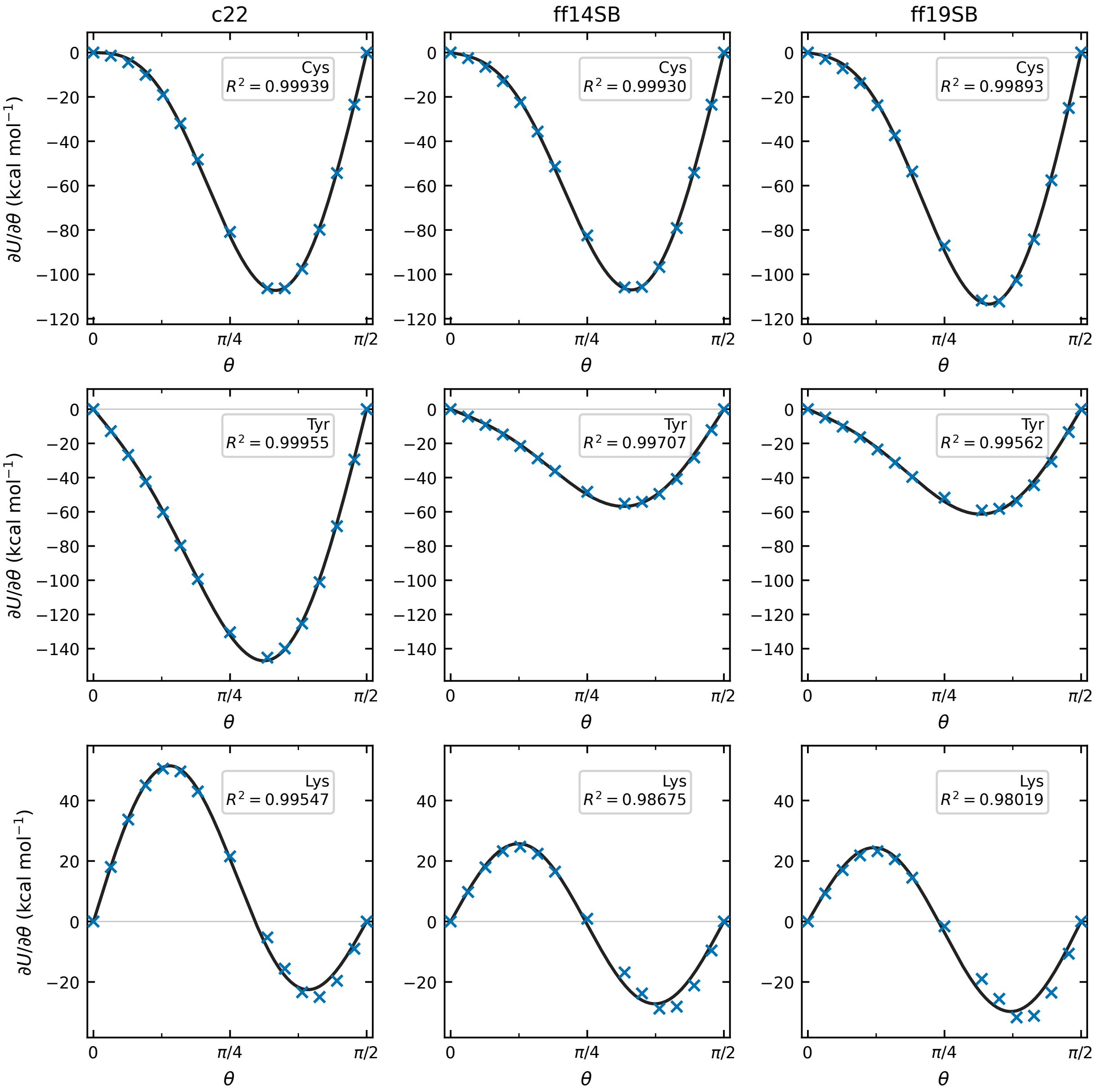

Figure S5: **Thermodynamic-integration fits for the Cys, Tyr, and Lys model compounds.** Rows correspond, from top to bottom, to Cys, Tyr, and Lys, and columns correspond, from left to right, to c22, ff14sb, and ff19sb. Each panel shows $\partial U/\partial\theta$ as a function of $\theta$. Symbols represent the generalized-force values used in the fits, and solid curves represent the fitted functions. Common vertical-axis limits are used within each row.

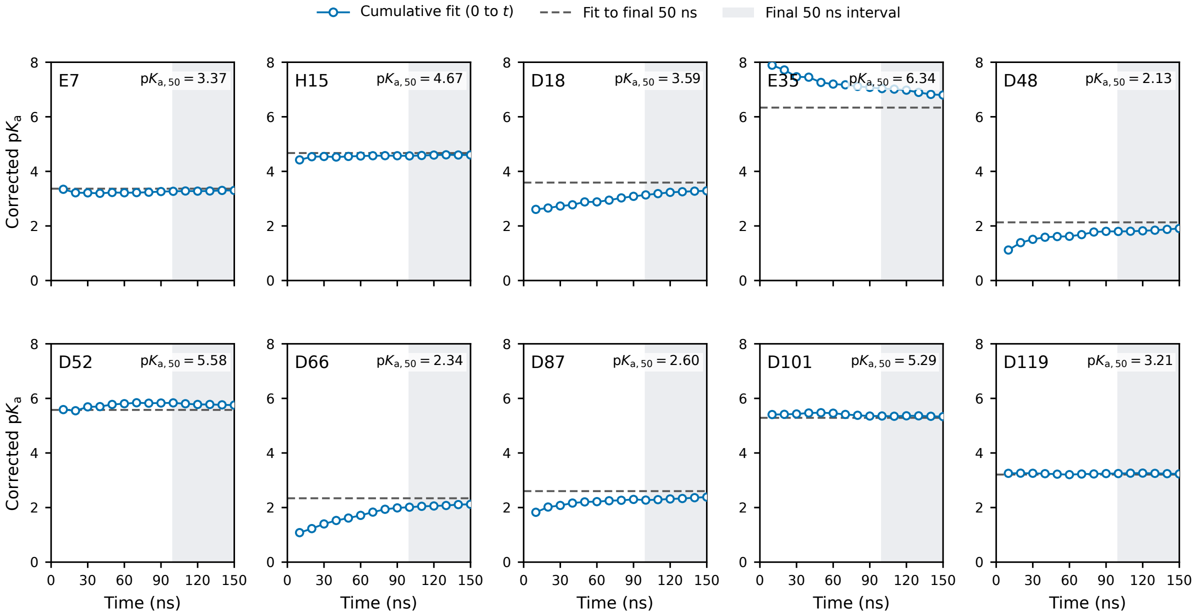

Figure S6: **Time evolution of calculated $\mathrm{p}K_\mathrm{a}$ values of HEWL in the pH-REX PME-CpHMD simulations with the c22 force field.** Cumulative $\mathrm{p}K_\mathrm{a}$ estimates were obtained at 10 ns intervals by fitting the unprotonated fractions across all pH conditions to the generalized Henderson–Hasselbalch equation followed by the finite-size correction. Dashed horizontal lines indicate the experimental $\mathrm{p}K_\mathrm{a}$'s. Shaded regions mark the final 50 ns using for $\mathrm{p}K_\mathrm{a}$ calculation and analysis in the main text (100–150 ns).

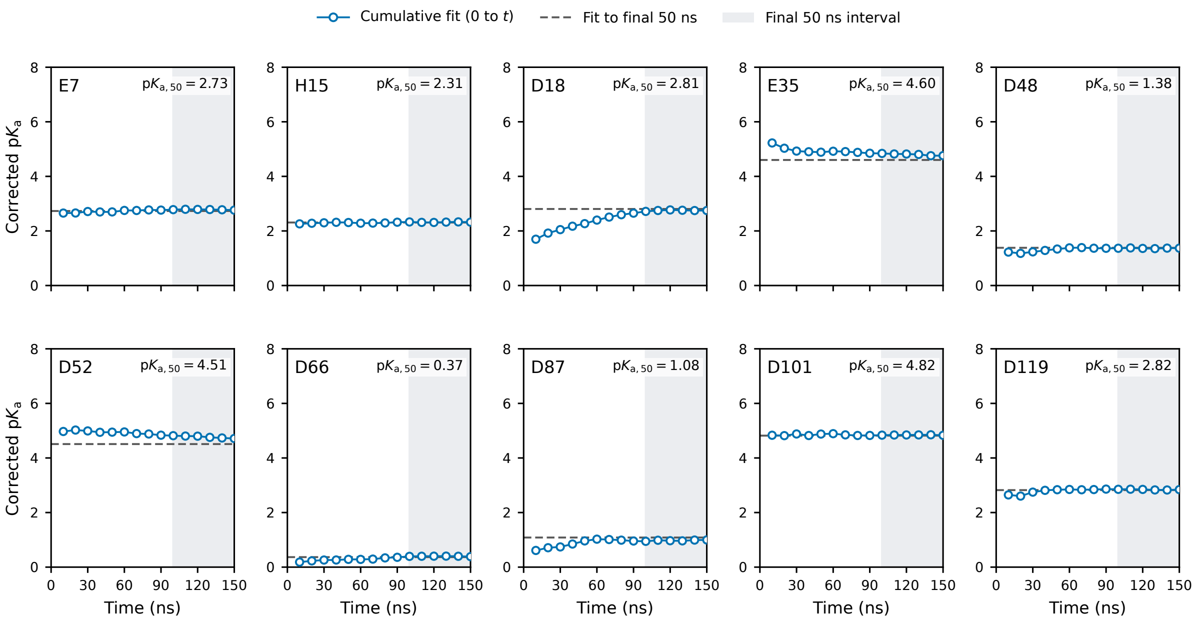

Figure S7: **Time evolution of calculated $\mathrm{p}K_\mathrm{a}$ values of HEWL in the pH-REX PME-CpHMD simulations with the ff14sb force field.** Cumulative $\mathrm{p}K_\mathrm{a}$ estimates were obtained at 10 ns intervals by fitting the unprotonated fractions across all pH conditions to the generalized Henderson–Hasselbalch equation followed by the finite-size correction. Dashed horizontal lines indicate the experimental $\mathrm{p}K_\mathrm{a}$'s. Shaded regions mark the final 50 ns using for $\mathrm{p}K_\mathrm{a}$ calculation and analysis in the main text (100–150 ns).

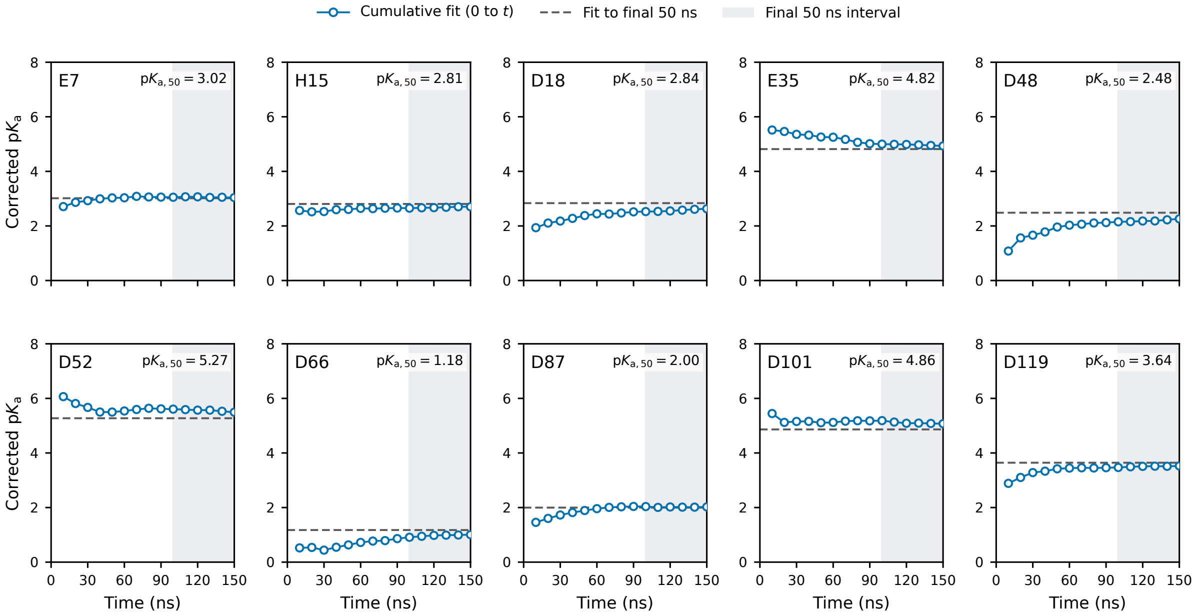

Figure S8: **Time evolution of calculated $\mathrm{p}K_\mathrm{a}$ values of HEWL in the pH-REX PME-CpHMD simulations with the ff19sb force field.** Cumulative $\mathrm{p}K_\mathrm{a}$ estimates were obtained at 10 ns intervals by fitting the unprotonated fractions across all pH conditions to the generalized Henderson–Hasselbalch equation followed by the finite-size correction. Dashed horizontal lines indicate the experimental $\mathrm{p}K_\mathrm{a}$'s. Shaded regions mark the final 50 ns using for $\mathrm{p}K_\mathrm{a}$ calculation and analysis in the main text (100–150 ns).

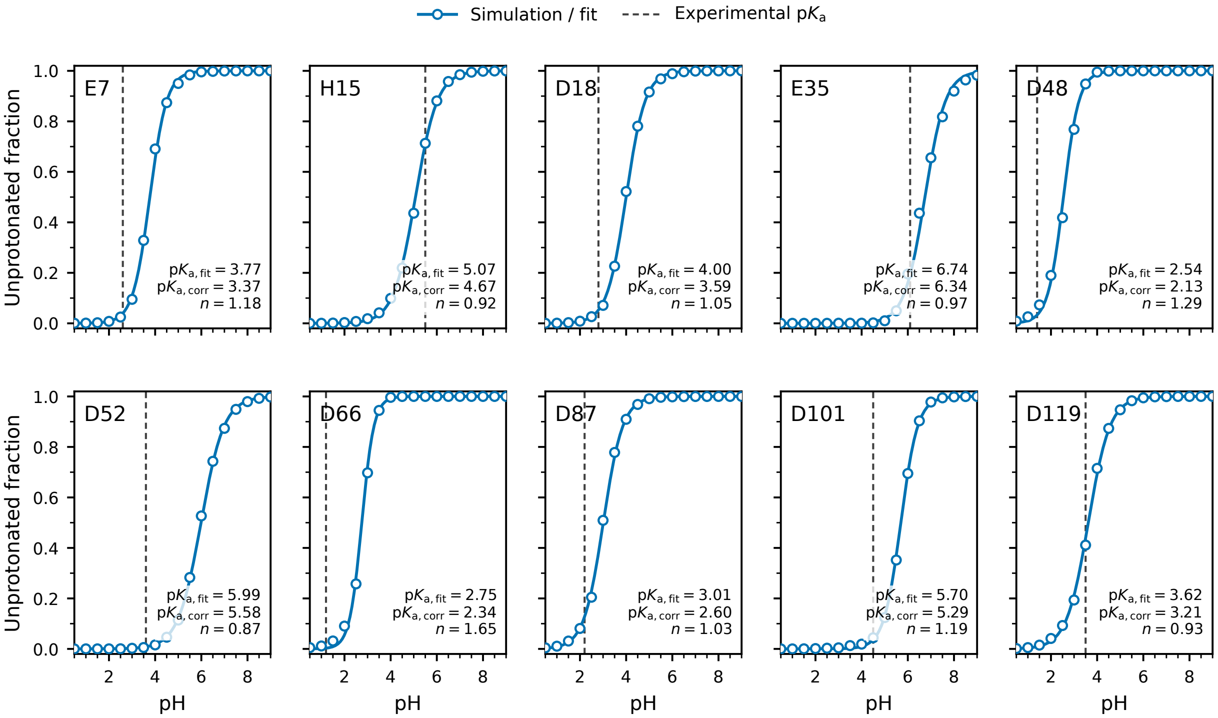

Figure S9: **Titration curves of HEWL residues from the c22 pH-REX PME-CpHMD simulations.** Symbols show the unprotonated fractions calculated over the final 50 ns at each pH condition, and solid curves show uncorrected fits to the generalized Henderson–Hasselbalch equation. Dashed vertical lines indicate the experimental $\mathrm{p}K_\mathrm{a}$ values. $\mathrm{p}K_\mathrm{a,fit}$ and $n$ denote the fitted $\mathrm{p}K_\mathrm{a}$ value and the Hill coefficient, respectively. $\mathrm{p}K_\mathrm{a,corr}$ denotes the $\mathrm{p}K_\mathrm{a}$ after the finite-size correction.

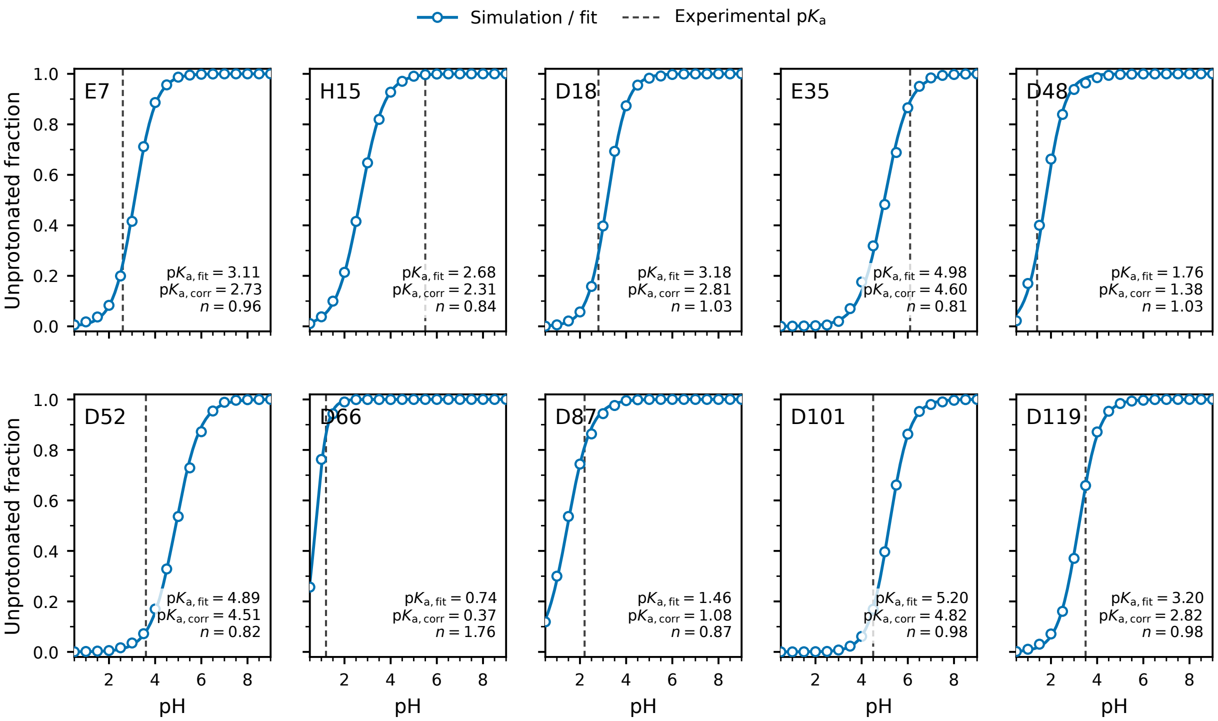

Figure S10: **Titration curves of HEWL residues from the ff14sb pH-REX PME-CpHMD simulations.** Symbols show the unprotonated fractions calculated over the final 50 ns at each pH condition, and solid curves show uncorrected fits to the generalized Henderson–Hasselbalch equation. Dashed vertical lines indicate the experimental $\mathrm{p}K_\mathrm{a}$ values. $\mathrm{p}K_\mathrm{a,fit}$ and $n$ denote the fitted $\mathrm{p}K_\mathrm{a}$ value and the Hill coefficient, respectively. $\mathrm{p}K_\mathrm{a,corr}$ denotes the $\mathrm{p}K_\mathrm{a}$ after the finite-size correction.

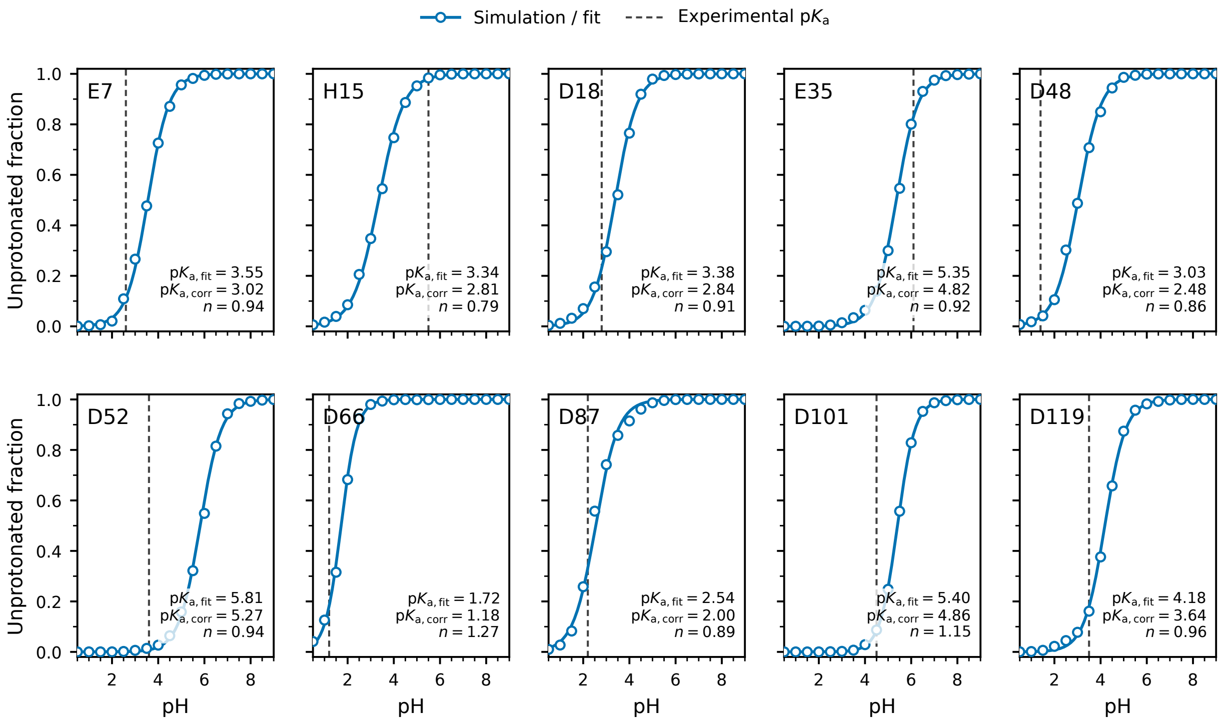

Figure S11: **Titration curves of HEWL residues from the ff19sb pH-REX PME-CpHMD simulations.** Symbols show the unprotonated fractions calculated over the final 50 ns at each pH condition, and solid curves show uncorrected fits to the generalized Henderson–Hasselbalch equation. Dashed vertical lines indicate the experimental $\mathrm{p}K_\mathrm{a}$ values. $\mathrm{p}K_\mathrm{a,fit}$ and $n$ denote the fitted $\mathrm{p}K_\mathrm{a}$ value and the Hill coefficient, respectively. $\mathrm{p}K_\mathrm{a,corr}$ denotes the $\mathrm{p}K_\mathrm{a}$ after the finite-size correction.

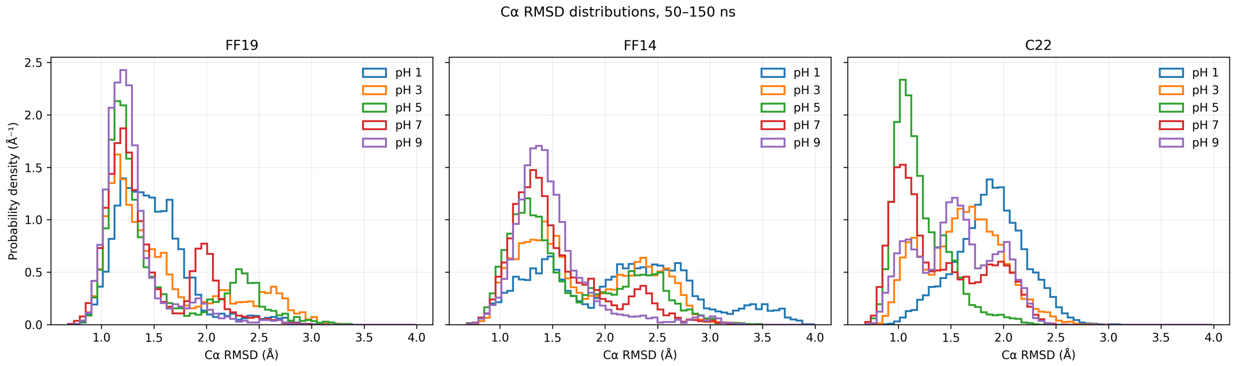

Figure S12: **C$\alpha$ RMSD distributions of hen egg white lysozyme (HEWL) from PME-CpHMD simulations with different force fields.** Probability density distributions at pH 1, 3, 5, 7, and 9 over 50–150 ns/replica simulations with the ff19sb (left), ff14sb (middle), and c22 (right) force fields. RMSDs were calculated using C$\alpha$ atoms for alignment and measurement, relative to the first analyzed frame of each pH trajectory.

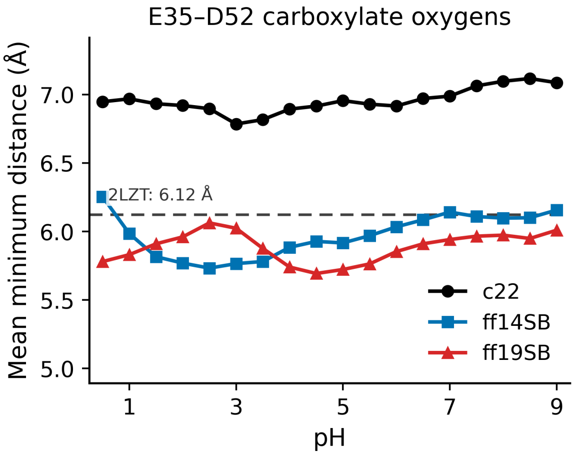

Figure S13: **Carboxylate oxygen separation between Glu35 and Asp52 in HEWL.** Mean minimum distance between the side-chain carboxylate oxygen atoms of Glu35 and Asp52 as a function of pH. The dashed horizontal line indicates the corresponding distance in the crystal structure (PDB ID: 2LZT).

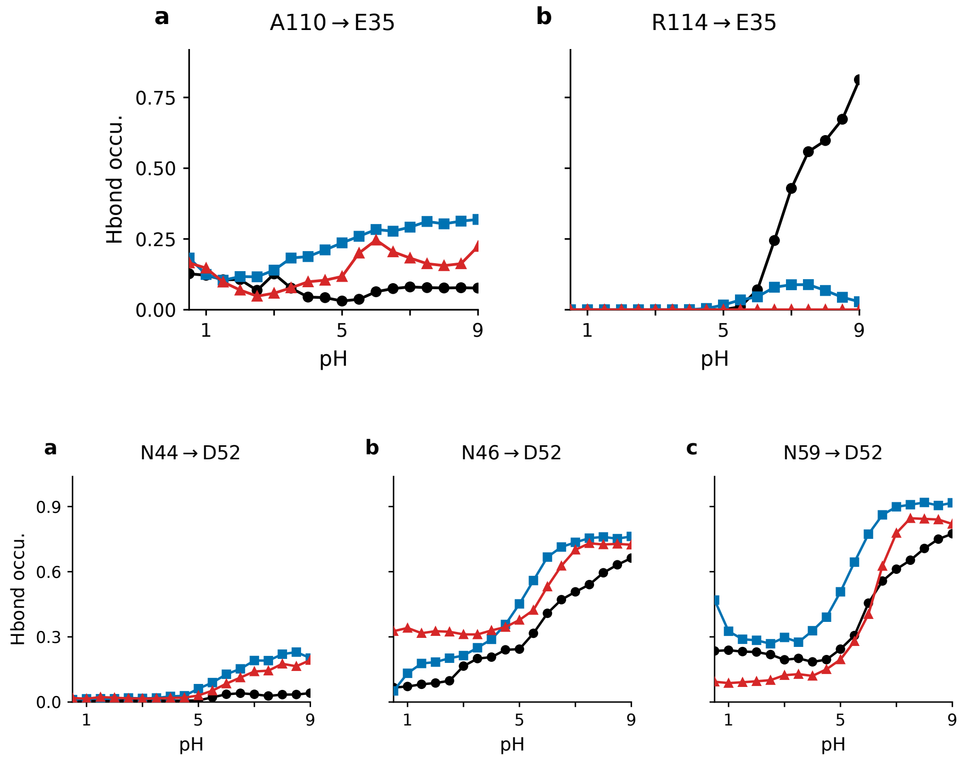

Figure S14: **Residue-specific hydrogen-bond occupancies of E35 and D52 in HEWL. Top panel.** pH-dependent occupancies of the h-bonds A110→E35 (**a**) and R114→E35 (**b**) from the c22 (black circles), ff14sb (blue squares), and ff19sb (red triangles) simulations. Occupancies represent the mean number of hydrogen bonds per frame. Their sum gives the E35 h-bond occupancy in Figure 4d of the main text. **Bottom panel.** pH-dependent occupancies of the h-bonds N44→D52 (**a**), N46→D52 (**b**), and N59→D52 (**c**) from the c22 (black circles), ff14sb (blue squares), and ff19sb (red triangles) simulations. Their sum gives the D52 occupancy in Figure 4d of the main text.

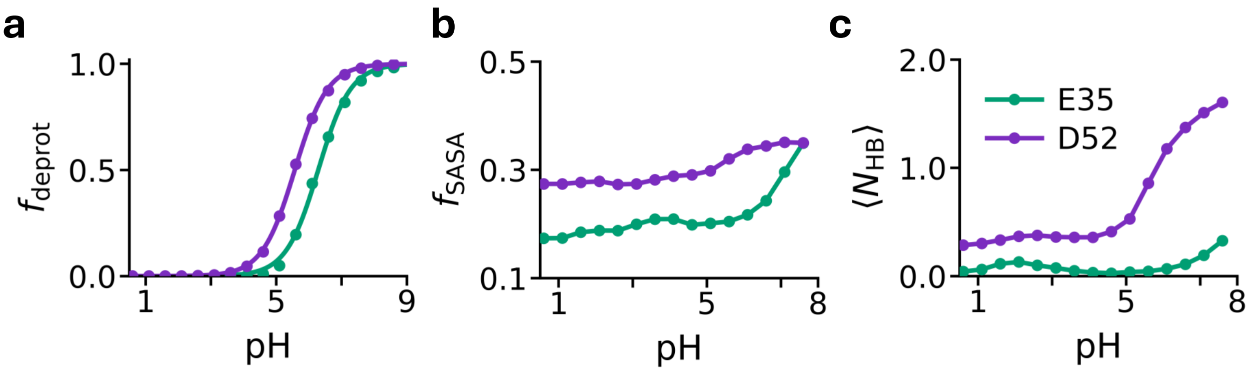

Figure S15: **Titration and local environments of E35 and D52 in HEWL from the PME-CpHMD simulations with the c22 force field at 50 mM ionic strength.** E35 and D52 are shown in green and violet, respectively. **a.** pH-dependent deprotonated fractions ($f_{\text{deprot}}$) of E35 and D52. Symbols represent simulation data, and solid curves are best fits to the generalized Henderson–Hasselbalch equation. **b.** pH-dependent fractional solvent-accessible surface area ($f_{\text{SASA}}$) of the carboxylate oxygens of E35 and D52, normalized by the corresponding model tripeptide values in solution. **c.** Mean number of hydrogen bonds ($\langle N_{\text{HB}} \rangle$) accepted by E35 from A110 and R114, and by D52 from N44, N46, and N59, as a function of pH. Hydrogen-bond counts are averaged over all analyzed frames, including frames with no hydrogen bonds. Data were analyzed over 3–153 ns of each pH trajectory. For all three panels, a pH correction of $-0.40$ was applied to the plotted pH coordinates, consistent with Fig. 4; the calculated observables were not altered.

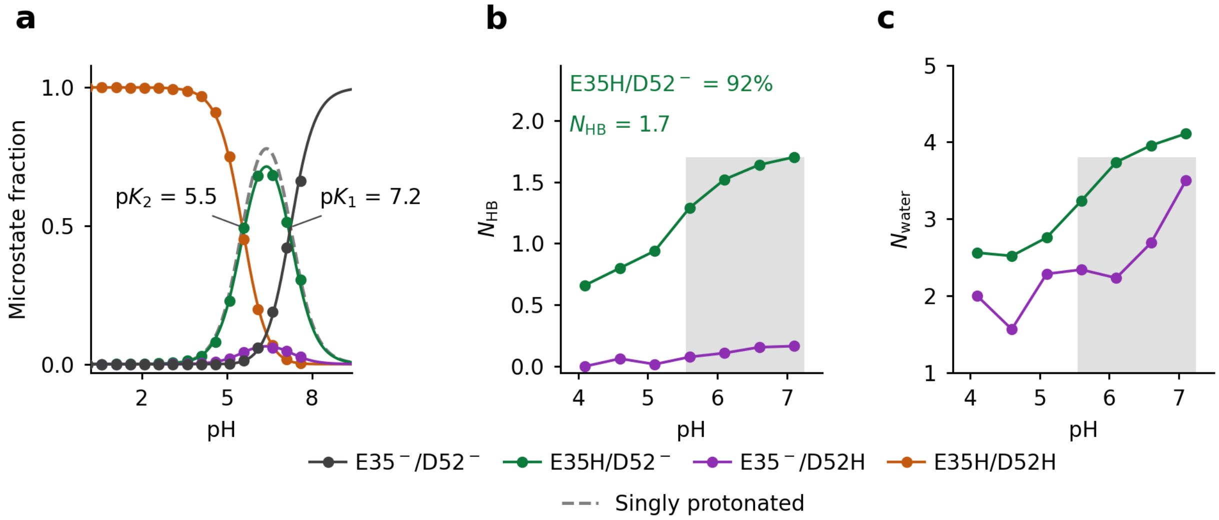

Figure S16: **Coupled titration and electrostatic environment of the singly protonated states of E35/D52 in HEWL from PME-CpHMD simulations with the c22 force field at 50 mM ionic strength. a.** pH-dependent fraction of the four microstate populations, E35H/D52H, E35H/D52$^-$, E35$^-$/D52H, and E35$^-$/D52$^-$. Curves are best fits to the four-state model. The dashed grey curve is the total population of the two singly protonated states; its crossings with E35$^-$/D52$^-$ and E35H/D52H give p$K_1$ and p$K_2$, respectively, obtained from fitting to the two-proton model. **b.** Number of h-bonds ($N_{\mathrm{HB}}$) accepted by the deprotonated carboxylate in E35$^-$/D52H (purple) or E35H/D52$^-$ (green). The percentage indicates the fraction of E35H/D52$^-$ within the singly protonated population. The annotated $N_{\mathrm{HB}}$ is the mean number of h-bonds for E35H/D52$^-$ (green) at p$K_1$. **c.** Number of water molecules ($N_{\mathrm{water}}$) within 3.4 Å of the deprotonated carboxylate oxygens (first-shell) in each singly protonated state. In panels **b** and **c**, the shaded region spans p$K_2$ to p$K_1$. All pH values include the $-0.40$ finite-size correction. H-bond is defined by a donor–acceptor distance $\leq$ 3.5 Å and an angle $\geq$ 150°.
